# Optimal spatial prioritization of control resources for elimination of invasive species under demographic uncertainty

**DOI:** 10.1101/812305

**Authors:** Kim M. Pepin, Timothy J. Smyser, Amy J. Davis, Ryan S. Miller, Sophie McKee, Kurt C. VerCauteren, Chris Slootmaker

## Abstract

Populations of invasive species often spread heterogeneously across a landscape, consisting of local populations that cluster in space but are connected by dispersal. A fundamental dilemma for invasive species control is how to optimally allocate limited fiscal resources across local populations. Theoretical work based on perfect knowledge of demographic connectivity suggests that targeting local populations from which migrants originate (sources) can be optimal. However, demographic processes such as abundance and dispersal can be highly uncertain, and the relationship between local population density and damage costs (damage function) is rarely known. We used a metapopulation model to understand how observational uncertainty in abundance and connectivity, and imperfect knowledge of the damage function, affect return on investment (ROI) for optimal control strategies. Budget, observational uncertainty, and the damage function had strong effects on the optimal resource allocation strategy. Uncertainty in dispersal probability was the least important determinant of ROI. The damage function determined which resource prioritization strategy was optimal when connectivity was symmetric but not when it was asymmetric. When connectivity was asymmetric, prioritizing source populations had a higher ROI than allocating effort equally across local populations, regardless of the damage function, but uncertainty in connectivity structure and abundance reduced ROI of the optimal prioritization strategy by 57% on average depending on the control budget. With low budgets (monthly removal rate of 6.7% of population), there was little advantage to prioritizing resources, especially when connectivity was high or symmetric, and observational uncertainty had only minor effects on ROI. Allotting funding for improved monitoring appeared to be most important when budgets were moderate (monthly removal of 13-20% of the population). Our result showed that multiple sources of observational uncertainty should be considered concurrently for optimizing ROI. Accurate estimates of connectivity direction and abundance were more important than accurate estimates of dispersal rates. Developing cost-effective surveillance methods to reduce observational uncertainties, and quantitative frameworks for determining how resources should be spatially apportioned to multiple monitoring and control activities are important and challenging future directions for optimizing ROI for invasive species control programs.

## INTRODUCTION

The interaction of reproduction and dispersal processes determines the dynamics and persistence of spatially-structured populations (Hanski 1998). Reproduction provides new individuals that can disperse among local populations, promoting persistence in the global population. Rates and directions of dispersal in space define the ‘demographic connectivity’ of a population. The concept of demographic connectivity underlies modern theory in conservation planning (Rudnick et al. 2012, van Nouhuys 2016). Conserving patches of habitat with higher levels of connectivity facilitates dispersal among local populations (Williams et al. 2004, Rudnick et al. 2012), which maximizes population growth rates and makes the entire population more resilient to extinction pressures (Hanski and Ovaskainen 2000, Williams et al. 2004, Hastings and Botsford 2006, Jacobi and Jonsson 2011).

An understanding of the importance of demographic connectivity is also used to optimize control of invasive species (Travis and Park 2004, Chades et al. 2011, Caplat et al. 2012, Glen et al. 2013, Baker 2017) and disease (Haydon et al. 2006, Chades et al. 2011). When control in spatially-structured populations of invasive species is the objective, general guidelines for optimal prioritization have included to ‘target the most highly-connected local populations’ (Perry et al. 2017) and to ‘target local source populations’ (Baker 2017) to minimize abundance and maximize extinction rates. These recommendations are consistent when considering a variety of connectivity structures, for example: managing upstream local populations when connectivity is directional, managing any local population and then its nearest neighbors when connectivity is a ring structure, and to start at one end and continue sequentially in bidirectional or linear connectivity structures (Chades et al. 2011). However, more complex recommendations can emerge when one local population is highly connected and all others have only one connection; i.e., ring structures (Chades et al. 2011), when local populations exhibit source-sink dynamics (Travis and Park 2004), and depending on population growth (Caplat et al. 2014) and dispersal rates (Travis and Park 2004). Thus, the optimal strategy is affected by context-dependent demographic processes, including population growth rates and dispersal dynamics.

In practice, using knowledge of spatial population processes to manage populations (‘connectivity-based management’) requires estimates of demographic connectivity among local populations. Measures of demographic connectivity can be based on landscape features, or on estimates of dispersal behavior or gene flow between local populations (Baguette et al. 2012). Landscape-based estimates (e.g., least cost path analysis) are useful for examining how processes such as landscape fragmentation and climate change might affect population connectivity and persistence, but usually neglect individual-level dispersal behavior (i.e., how animals actually travel through the landscape) (Baguette et al. 2012). Estimates from tracking individual movement over time are ideal for understanding and predicting population persistence, but directly monitoring dispersal behavior for many individuals can be expensive in management applications. To address this gap, measures of gene flow have been used to infer demographic connectivity (Lowe and Allendorf 2010). However, models for estimating gene flow between local populations can require unrealistic or inappropriate assumptions (e.g., genetic equilibrium conditions, narrow ranges of demographic conditions), resulting in biased or imprecise estimates (Faubet et al. 2007, Meirmans 2014, Younger et al. 2017). Also, genetic estimates of demographic connectivity can be highly uncertain (Samarasin et al. 2017). Thus, the practical benefit of using estimates of connectivity based on real-world data to inform optimal allocation of control resources remains poorly understood.

Applying optimal resource allocation strategies through connectivity-based management requires monitoring data on abundance, demographic connectivity, and the damage function. In addition to structural uncertainty in the ecological and management processes that determine the true levels of abundance, the data used by managers to implement connectivity-based management are uncertain due to observational error. Observational uncertainty in monitoring data of invader presence or abundance can make theoretically optimal strategies less effective (Chades et al. 2011, Regan et al. 2011, Rout et al. 2014, Kling et al. 2017, Bonneau et al. 2019). Similarly, as the optimal resource allocation strategy depends on the relationship between invader abundance and damage costs (Yokomizo et al. 2009, Davis et al. 2018a), imperfect knowledge in this relationship could affect the optimal strategy. And, yet to be determined, are the potential effects of imperfect knowledge about the true connectivity structure.

Another major gap is that the relative importance of these different sources of observational uncertainty on optimal resource allocation decisions to control invasions remains unexplored. To address this gap, we built on work that separately examines how observational uncertainty in abundance (Haight and Polasky 2010, Kling et al. 2017) and knowledge of the damage function (Yokomizo et al. 2009, Davis et al. 2018a) affects the optimal resource allocation strategy by concurrently considering these effects with those from observational uncertainty in connectivity in a single framework. Specifically, our objective was to provide guidance for using one-time connectivity estimates to inform surveillance and control decisions in practical settings. We predicted that observational uncertainty in the one-time estimate of connectivity and regular estimates of abundance would decrease the benefits of prioritizing source populations because there would be increased error in prioritizing resources among local populations. We also expected that the optimal resource allocation strategy would depend on knowledge of the damage function. Our results provide general guidance for spatial prioritization of resources under multiple sources of uncertainty and fixed budgets and highlight which data may be more valuable to collect for optimizing resource allocation.

## METHODS

### Invasive species and management context

Our theoretical framework is inspired by demographic dynamics of invasive wild pigs (*Sus scrofa*) in areas of the U.S.A. where populations of wild pigs are not contiguous; although our approach was developed to apply more generally to the control of any spatially-structured pest population. In areas of the U.S.A where wild pig populations are less abundant and along the edges of areas with high-density pig populations, their distribution is similar to a metapopulation with distinct subpopulations and genetic evidence of recent dispersal between subpopulations (Smyser, T., unpublished data). Wild pigs produce offspring throughout the year (Mayer and Brisbin 2009), with 1-2 seasonal peaks in parturition rather than 1 distinct seasonal birth pulse. A genetic archive of wild pig samples collected ancillary to ongoing control efforts is currently being developed to guide management decisions based on spatial dispersal probabilities and directions. We used a range of probabilities and simplified connectivity structures that have been determined for wild pigs from genetic analyses (Tabak et al. 2017, Hernandez et al. 2018), Smyser, T. unpublished data) to determine optimal allocation of resources across populations where the management objective is to eliminate the metapopulation at the lowest overall cost of damages and removal costs.

### Overview of approach

Similar to previous work (Chades et al. 2011, Baker and Bode 2016, Baker 2017), our framework accounts for non-linear relationships between population density and removal and damage costs, in determining the optimal resource allocation strategy. However, we simplify the number of potential resource allocation strategies *a priori* by considering practical rules of thumb (‘prioritization schemes’) that could be implemented by managers at coarse time scales. Thus, our method aims to identify the best strategy for the objective function (defined below) within a pre-defined set of strategies (hereafter referred to as ‘optimal’), rather than the optimal solution from all possible strategies (i.e., we conducted simulation optimization (Amaran et al. 2014, Punt et al. 2016)). We define the optimal resource allocation strategy as the one with the highest return on investment (ROI; ‘returns’, *i.e*., savings from management, divided by its management costs – see Eq. 14 below).

We use a stochastic metapopulation model to allow for natural levels of uncertainty (stochasticity) that would be expected in the true management (number of individuals removed relative to the intended number) and ecological (abundance in each subpopulation) processes. These sources of uncertainty are structural – affecting the generation of the true abundance. To this we add observational uncertainty in abundance and connectivity. We assume that the manager has an estimate of connectivity among local populations through a genetic analysis (or other method) conducted at the outset of a new management program. We also assume that the manager has monitoring data on population abundance at each time step through a method that requires no additional cost, e.g., analyzing the removal data using a removal model (Davis et al. 2016). We first examine how applying different prioritization schemes to different connectivity structures and damage functions affects ROI when abundance and connectivity are perfectly observed. Then, we examine how observational uncertainty in the one-time estimate of connectivity and the monthly estimates of abundance affect ROI of the assumed optimal strategy. Our approach is conceptually similar to bioeconomic models such as Haight and Polasky (2010) and Regan et al. (2011) in that we examine the effects of observational uncertainty on optimal resource allocation strategies. However, our approach differs from these approaches in two important ways: 1) we assume that the manager pre-determines how resources should be allotted in space based on knowledge of connectivity structure and test how observational uncertainty in connectivity and abundance monitoring affects ROI of an assumed optimal resource allocation strategy, rather than determining how much should be apportioned to monitoring throughout control in order to apply the optimal strategy, and 2) our ‘strategies’ are defined by how a fixed amount of resources are apportioned in space rather than how they are apportioned to control versus monitoring. In our approach, we assume all monitoring data are available at no additional cost.

Code for the simulation model is given in Data S1. The model processes at each time step progress as follows: 1) density-dependent growth of each subpopulation (Eq. 1, 2), 2) dispersal among subpopulations according to their connectivity structure and rates (Eq. 3, 4), 3) management removal of individuals from each subpopulation (Eq. 5) dependent on the abundance-capture success relationship (Eq. 6) and the effort allotted to each subpopulation which depends on the overall effort available (*β*) and an effort prioritization function (*f_p_*) among subpopulations (Eq. 7, 8), 4) determination of management costs dependent on multiplying overall effort by a dollar value (Eq. 9), and, finally, 5) determination of the damage value dependent on density (Eq. 10) and on the shape of the density-damage relationship that is fixed throughout each simulation (either Eq. 11, 12, or 13). Lastly, ROI is calculated for each simulation from the cumulative management costs, cumulative damage costs, and elimination status at the end (Eq. 14). Each of these processes are described in detail in the following.

### Metapopulation model

We consider an established metapopulation of an invasive species that is spatially-structured with three subpopulations connected through dispersal, and closed to emigration and immigration to and from outside sources. We also assume that each subpopulation has a fixed carrying capacity (i.e., pre-defined geographic capacity) such that subpopulations cannot expand outside their patches. We examine 6 different connectivity ‘motifs’ within the metapopulation to study the effects of connectivity structure on the optimal management strategy (Fig. 1). We assume that each subpopulation can expand locally (up to their carrying capacity) and that subpopulations can be completely removed, but that no new subpopulations can be added. At the start of control, we assume that the metapopulation is at carrying capacity (i.e., equal abundance of subpopulations) and that control efforts decrease metapopulation abundance below carrying capacity.

**Fig. 1.**
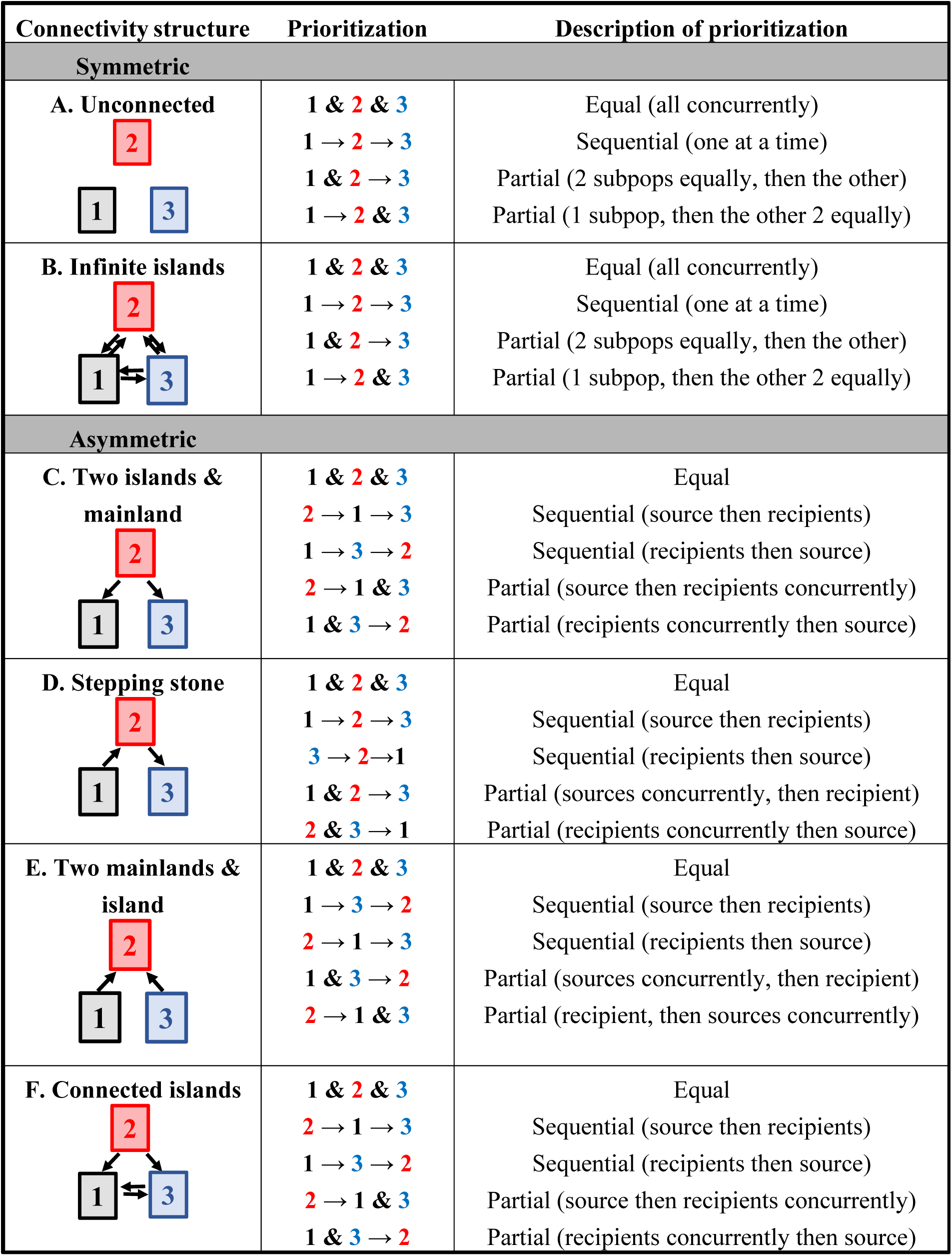
Connectivity motifs and a description of prioritization schemes. Connectivity increases with the number of arrows.

### Reproduction

We assume subpopulations in the metapopulation undergo logistic population growth to a fixed carrying capacity. Birth dynamics occur in discrete time at a monthly time step with Poisson-distributed error.

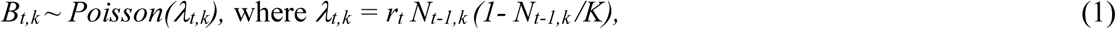

with abundance dynamics:

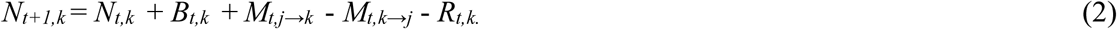

Here *B_t,k_* are the new births in month *t* in subpopulation *k,* and *N_t,k_* is population abundance. *M_j→k_*, *M_k→j_,* and *R_t,k_* (described in detail below) are the number of immigrants from each subpopulation *j* to subpopulation *k*, the number of emigrants from subpopulation *k* to each subpopulation *j*, and the number of invaders removed due to control. We assume the fixed carrying capacity, *K*, is equal to 1000 in each subpopulation and that abundance can exceed this level slightly due to stochastic processes. *r_t_* is a monthly growth rate that fluctuates based on seasonal birth dynamics (Table 1). Each subpopulation is initialized at carrying capacity.

**Table 1.**
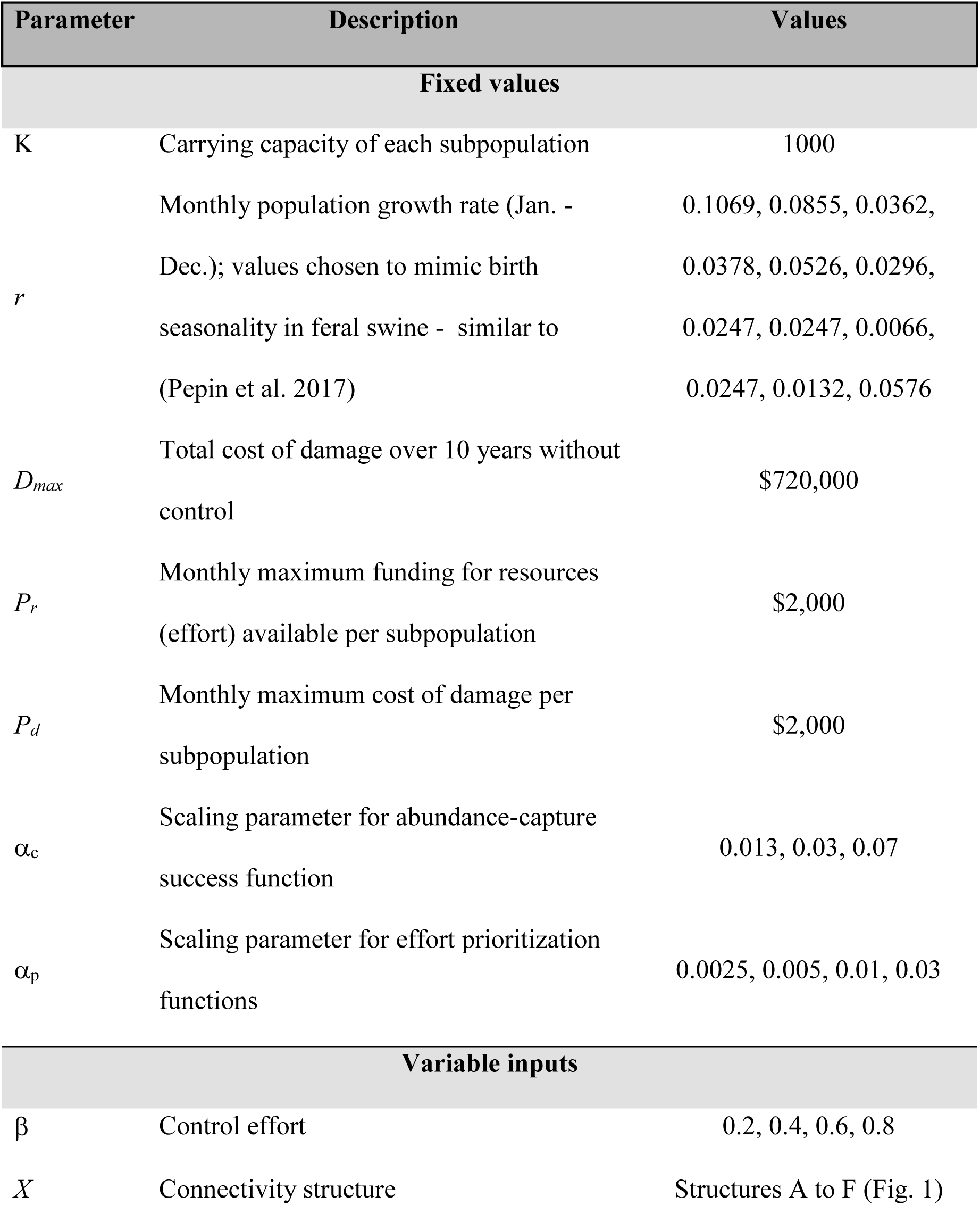
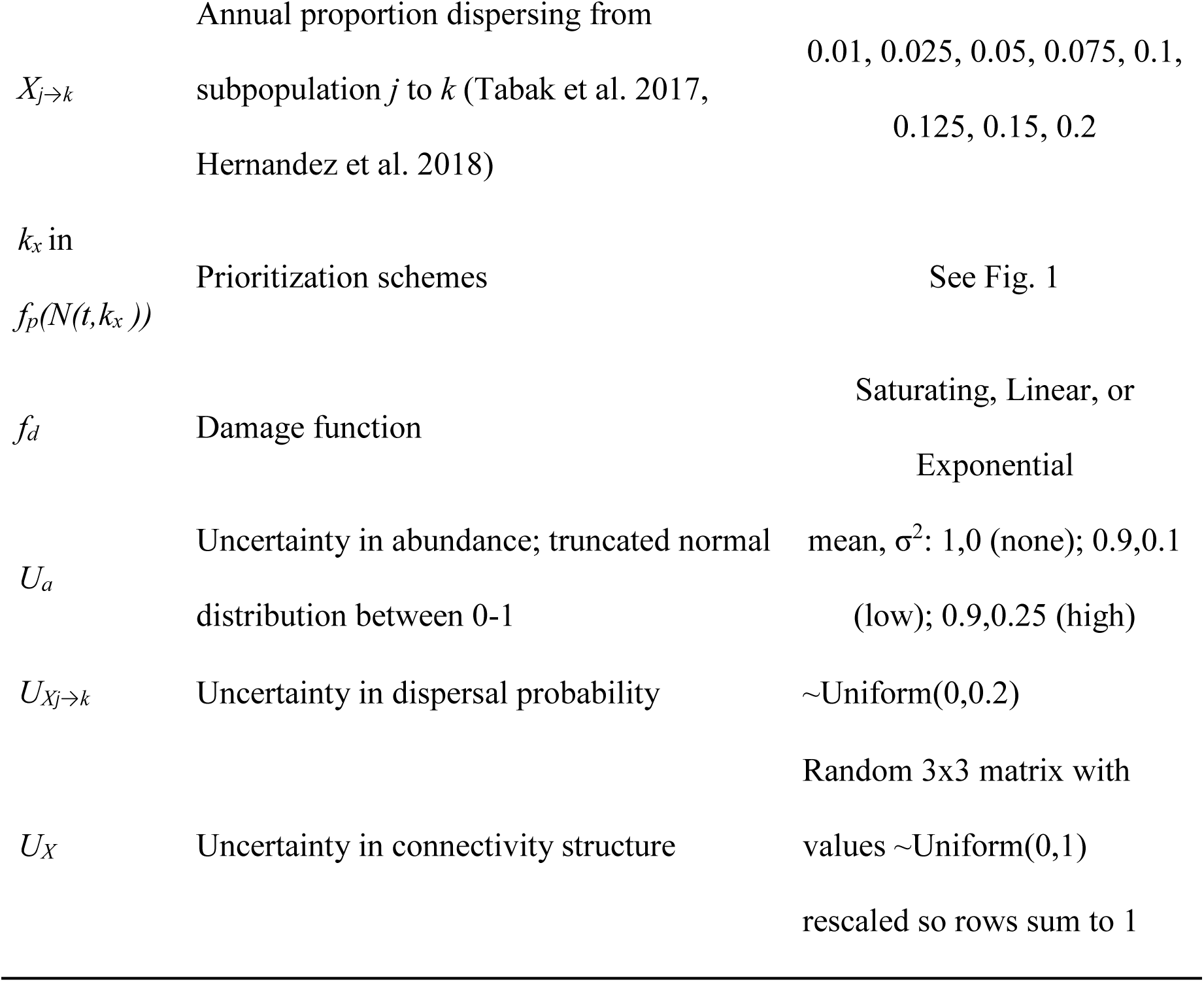
Description of parameters.

### Dispersal

Subpopulations are potentially connected by dispersal. We consider 6 different dispersal structures: A) Unconnected, B) Two islands and mainland, C) Connected islands, D) Two mainlands and island, E) Infinite islands, and F) Stepping stone (Fig. 1). At each time step, new migrants are added to subpopulation *k* as follows:

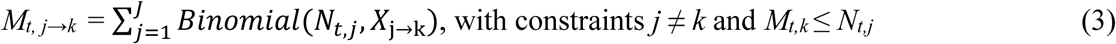

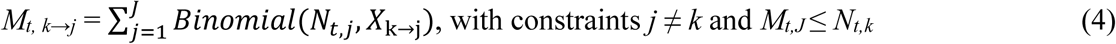

*X*_j→k_ is the monthly dispersal probability from subpopulation *j* to *k*. Thus, we assume that dispersal can occur in any month, depending on fixed dispersal probabilities among subpopulations, and that dispersal probabilities are proportional to abundance (i.e., not density-dependent).

### Control

In the absence of prioritization, a fixed monthly control effort (*β*) is always divided equally among subpopulations. We assume that the number of individuals removed from a population *k* in month *t*, *R_t,k_*, is a random variable that depends on effort allocated, and declines steeply with the population size as individuals become hard to find (Choquenot et al. 1999). We model R as a Poisson random variable:

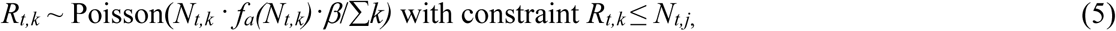

where *f_a_* is the abundance-capture success function and ∑*k* is the total number of subpopulations. The *f_a_* function quantifies what proportion of the population is available to be removed at each time step. We follow the intuition of (Choquenot et al. 1999), defining *f_a_* as Eq. 6.

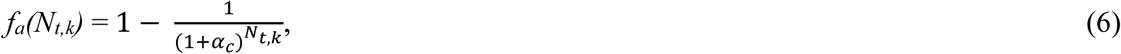

*f_a_* assumes that the proportion of the population that can be removed using a fixed amount of effort increases sharply at lower abundances to an asymptote of 1 when abundance is high because individuals are more easily located at high abundance and thus removal success (the proportion of the target removal number) increases to full capacity (i.e., 1). For example, if α_c_ = 0.03 and abundance is > 200, then the target removal number will almost always be achieved. However, the actual number removed will be lower than the target removal number when abundance is < 200 pigs with α_c_ = 0.03. α_c_ is a tuning parameter affecting the threshold abundance where actual management removal becomes lower than the target number (Appendix 1: Fig. S1). In general, lower values of α_c_ lead to less efficient removal (lower success), whereas higher values lead to more efficient removal (higher success). By modeling the effects of removal using a Poisson variable, we allow for stochastic variation in abundance to scale to mean abundance.

When prioritization of subpopulations occurs, the fixed monthly control effort, *β,* is not divided equally rather it is apportioned to subpopulations using a prioritization function (*f_p_*). We assume that the manager has abundance estimates at each time step to determine the amount of effort that should be allotted to each subpopulation based on *f_p_*, and that effort would begin to be shifted from the most prioritized subpopulation to less prioritized subpopulations as abundance decreases because fewer locations within a subpopulation would need to be targeted at lower abundance. Thus, in general, when prioritization occurs, the number of individuals removed at time *t* in subpopulation *k* is:

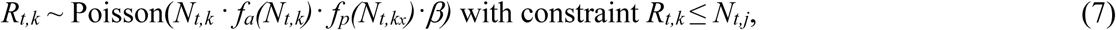

where *f_p_(N_t,kx_)* is conditional on the spatial prioritization scheme (*k_x_*, Fig. 1) for removal and current *N*_t,k_ and α_p_ according to:

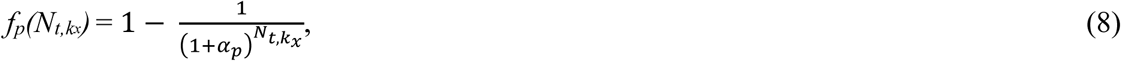

where α_p_ scales the amount of effort that will be allotted to subpopulation *k_x_* (specifies the prioritization scheme) as a function of its abundance. Lower values of α_p_ generate prioritization rules that are closer to equal allocation whereas higher values lead to higher discrepancy in the amount of effort allotted to each subpopulation (Appendix 1: Fig. S2). Appendix 1: Fig. S2 shows the effects of α_p_ on example patterns of effort allotment to the most prioritized subpopulation (*k_x=_*_1_), followed by secondarily and lastly prioritized subpopulations. In addition to equally dividing resources (as above), we allow three potential prioritization schemes (*k_x_*): 1) sequential (*k_x=_*_1 then 2 then 3_), 2) first two equally and then the last (*k_x=_*_1 & 2 equally, then 3_), and 3) first one and then the other two equally (*k_x=_*_1 then 2 & 3 equally_; Fig. 1). Specific equations defining how effort is allotted under different general prioritization schemes (1-3 above) and values of α_p_ are described in Data S1: culling3.

### Removal costs and damage function

We assume that there is a fixed monthly budget available to control the metapopulation with upper maximum *P_r_*. When effort is divided evenly among the subpopulations, the maximum monthly effort at high abundance in each subpopulation results in removal of 33.3% of each subpopulation monthly (i.e., if β = 1, which we do not examine). We use an arbitrary maximum monthly budget of $6000 for control of the metapopulation ($2000 per subpopulation when effort is divided equally; with all monetary amounts expressed in real dollars). Different levels of control effort (β = 0.8, 0.6, 0.4, 0.2) correspond to decreases in the monthly budget for control by 20, 40, 60, or 80%, which translates to maximum monthly removal rates of 26.6, 20.0, 13.3, and 6.7%, in each subpopulation when abundance is high and resources are divided equally. For example, under the lowest control effort (β = 0.2), the maximum amount of monthly funding available to control is $1,200 while the maximum amount of monthly damage at carrying capacity is $6,000. Thus, β determines both the proportion of individuals that can be removed monthly and the relative difference between damage and control costs. Also, as mentioned above, the same amount of resources become less efficient as population density declines such that removal success decreases with the same amount of effort once subpopulations reach low densities. Total monthly amount spent on control is as follows:

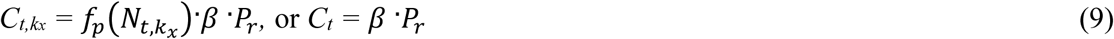

where *C_t,k_x* is the amount spent on subpopulation *k_x_* in month *t*, *C_t_* is the amount spent for all subpopulations in month *t*, and *P_r_* is the fixed maximum amount of resources available to be spent each month for removal ($6000).

We examine three different damage functions (*f_d_(N_t,k_)*) for determining damage costs based on abundance (Eq. 11-13). We assume that total monthly damage costs from the metapopulation (*D_t_*) at carrying capacity are a fixed arbitrary value ($6000 for the metapopulation; $2000 / subpopulation).

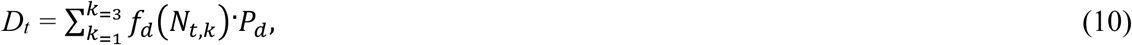

where *P_d_* is the maximum cost of damage for a subpopulation when it is at carrying capacity ($2000) and *f_d_* is the proportional damage as a function of current abundance. The different damage functions are described as follows:

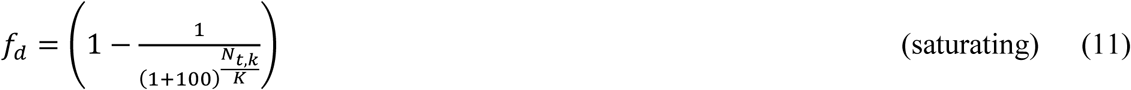

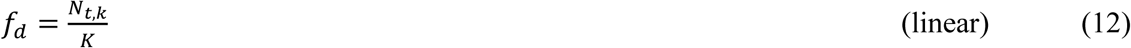

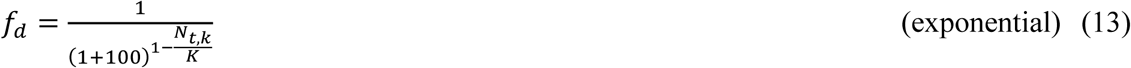

These damage functions are similar to Type II, Type I, and Type IV functional responses (Holling 1959), except our Type IV (exponential) function does not saturate at high abundance after rising from the low damage condition (Appendix 1: Fig. S3). We assumed the damage function was fixed throughout each simulation. We tested the effects of imperfect knowledge of the damage function by determining the optimal resource allocation strategy for each damage function.

### Evaluation

We evaluated outcomes of simulations by ROI, a measure of cost-effectiveness. ROI calculates the difference between total costs in the absence of control and total costs with control (i.e., damages avoided net of control costs) scaled to the cost of control. Thus, ROI gives the amount saved per dollar spent on control. ROI was calculated as:

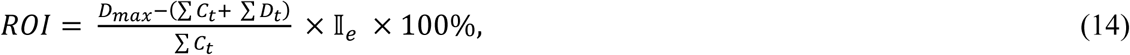

where *D_max_* is the total damage cost over 10 years when no control occurred, and I_e_ indicates whether the metapopulation was eliminated (I_e_ = 1) or not (I_e_ = 0) by the end of the simulation. Thus, we assumed that runs that did not eliminate had a return on investment of 0 because our objective was to identify strategies that led to elimination with the highest return on investment.

### Observational uncertainty in abundance

Our prioritization methods assumed that the manager has monitoring data that can be used to estimate abundance in the subpopulations at each time step (e.g., Davis et al. 2016). Because abundance measures are typically imperfect, we examined how uncertainty in the monthly estimates impact ROI of the optimal prioritization plan (Kendall and Moore 2012). We examined two levels of uncertainty in abundance (*U_o_*: low and high). We assumed that the process for observing abundance is biased low (underestimation of the true abundance) because studies that have estimated detection probability during management-based monitoring in populations have found that it is well below 1 (e.g., (Davis et al. 2018b), although in the supplementary information we also present results for instances where abundance estimation is imperfect but unbiased; Appendix 1: Fig. S10). We assumed the error term is a normally-distributed random variable truncated between 0 and 1 (i.e., biased low), with mean 0.9 and variance of 0.1 (low error) or 0.25 (high error). For each subpopulation at each time step, we multiplied the random proportion variable by the true abundance to determine the ‘observed’ abundance. Then, effort prioritization at each month was determined based only on the observed abundance and the prioritization function (*k_x_* scheme and α_p_).

### Observational uncertainty in connectivity

Because measures of connectivity can be difficult to obtain, we assumed that the manager had connectivity information once at the start of control (one-time information). We examined the situation where the manager assumes the one-time connectivity information is perfectly observed and remains the same over time, and thus applies the optimal control strategy in the case where abundance and connectivity are observed perfectly. To evaluate effects of uncertainty in connectivity structure, we apply the optimal strategy for each connectivity structure to itself and every other connectivity structure and calculate the mean and variance in ROI across all connectivity structures. Thus, our methodology evaluates how observing the wrong connectivity structure and thus potentially applying a sub-optimal allocation strategy affects the ROI outcome. Note that for some connectivity structures the same allocation strategy is optimal and thus observing connectivity structure imperfectly may not be problematic. We tested both aspects of imperfect knowledge in connectivity – the structure (pathways in Fig. 1) and the rates (dispersal probability among the edges). To evaluate effects of uncertainty in structure, we held the rate constant at 0.1 and evaluated ROI across all 6 structures under each optimal control strategy. To evaluate the effects of rates, we held the structure constant and evaluated ROI across all dispersal probabilities (Table 1). When evaluating uncertainty in both components we evaluated ROI for all structures and dispersal probabilities. In order to compare the uncertainties on equal grounds we calculated the means and their coefficients of variation by subsampling the data at random so that each metric was composed of 100 data points because there were more data points when both types of uncertainties were combined.

### Simulations

Each subpopulation was initialized with 1000 individuals and thus an equal level of damage. Each parameter set was run for 10 years of population dynamics and replicated 100 times. Design of parameter sets in simulations is given under ‘variable inputs’ in Table 1. To evaluate the effects of observational uncertainty on the optimal resource allocation strategy, we only ran the optimal spatial prioritization schemes (each optimal scheme ran on all 6 connectivity structures), and fixed *f_d_* at linear, *a_p_* at 0.03, and *a_c_* at 0.03, because effects of these parameters were examined in the first set of analyses (without uncertainty in abundance and connectivity). We did include variation in β in the uncertainty analyses because it was such a strong predictor of ROI. We performed all simulations and analyses in Matlab R2018a (MathWorks, Inc., Natick, MA). We evaluated how observational uncertainty in abundance and connectivity affected the optimal resource allocation strategies using: 1) descriptive summaries of mean ROI and the its uncertainty in terms of a coefficient of variation (reported below), 2) random forest analysis on the ROI outcomes to examine the relative importance of variables (methods described in caption of Appendix 1: Fig. S7), and 3) a zero-inflated negative binomial model on the ROI response to estimate the direction of effects of different variables (methods described in caption of Appendix 1: Table S1).

## RESULTS

### Control effort and damage function

Under the lowest β and no connectivity or prioritization (0.2; removal of 6.7% of the metapopulation monthly at carrying capacity; budget of $1200 monthly), control efforts failed to eliminate the metapopulation within 10 years, even when the abundance-capture success parameter (*α_c_*) was high (Appendix 1: Fig. S4), unless the subpopulations were sequentially prioritized (Appendix 1: Fig. S3C-F versus S4C-F). At a removal intensity of 0.4 (13.3% of the metapopulation monthly at carrying capacity; budget of $2400 monthly), when there was no prioritization complete elimination only occurred sometimes unless *α_c_* was high (Appendix 1: Fig. S4). However, if the subpopulations were sequentially prioritized, elimination occurred more often (ROI was higher) even when was *α_c_* moderate (Appendix 1: Fig. S5). A control effort of at least 0.6 (20% of the metapopulation monthly at carrying capacity; budget of $3600 monthly) and a moderate or high *α_c_* (0.03, 0.07) was required to eliminate the metapopulation frequently, with ROI being higher when removal was sequentially prioritized. (Appendix 1: Fig. S4, S5). Some form of prioritization almost always outperformed equal allocation of effort across the range of control effort values, except when the subpopulations had full, symmetric connectivity (Appendix 1: Fig. S6), but the magnitude of benefit from prioritization depended on control effort (e.g., Appendix 1: Fig. S6E). Relative to the effects of β, the effects of the damage function (*f_d_*) were more subtle (Fig. 2A versus 1B). ROI was generally higher when α_p_ was higher, meaning that more stringent prioritization (less like equal allocation) was generally better (Fig. 2C). Capture success (controlled by α_c_) had a very strong effect on ROI (Fig. 2D), where more efficient capture (higher values of α_c_) lead to much higher ROI.

**Fig. 2.**
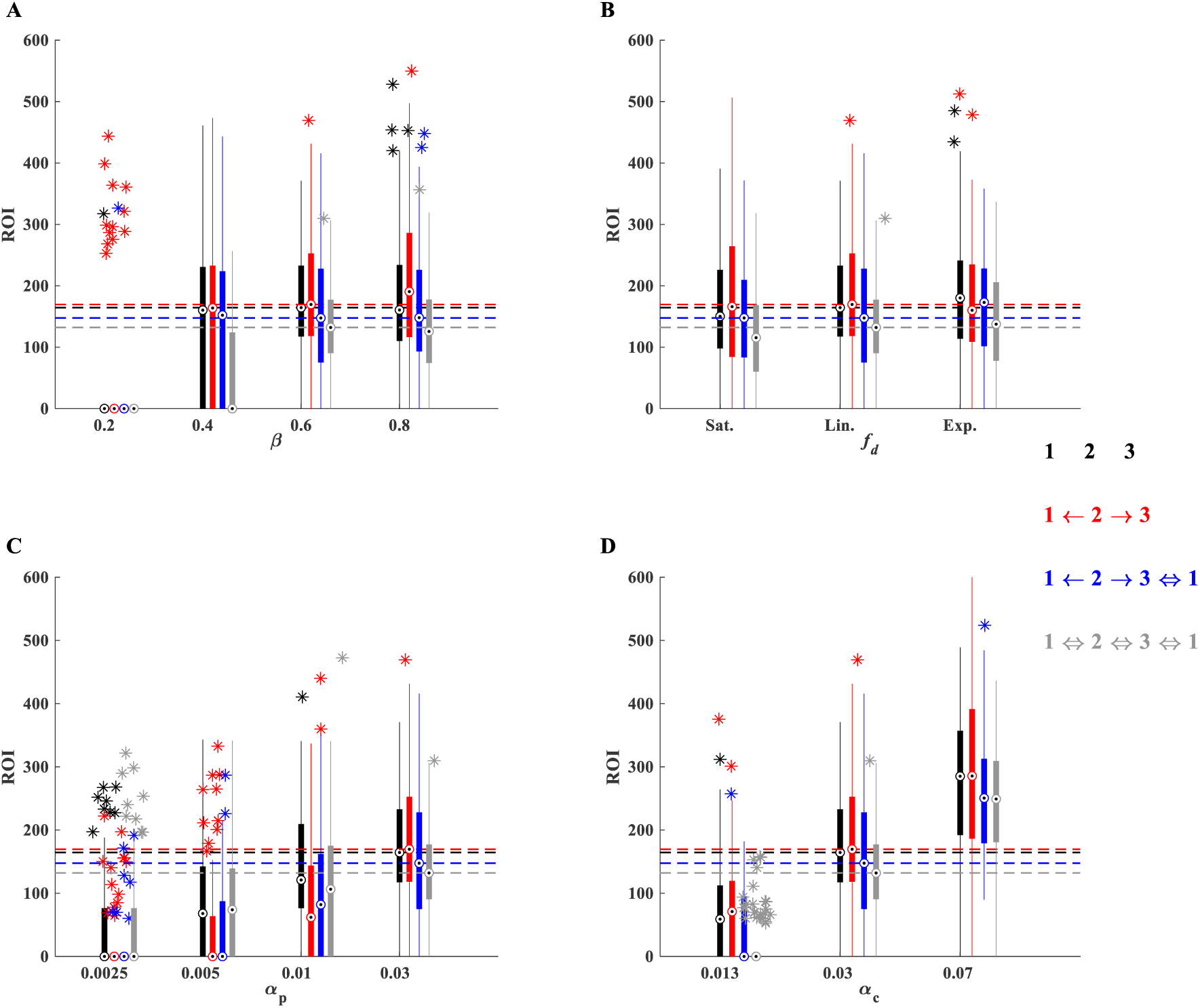
One-way sensitivity analysis of key parameters on ROI (%) for the optimal prioritization rule (shown as stars in Fig. 2). A) Effects of control intensity (budget); B) Damage function; C) Prioritization parameter, α_p_; D) Capture success parameter, α_c_. Boxplots summarize ROI for 100 replicate simulations. Each series of boxplots represents results from a different connectivity structure (see legend). Stars are outliers in the boxplots distributions. Horizontal dashed lines indicate the median ROI for the default parameter values (i.e., values that were fixed in Fig. 2; β = 0.6, *f_d_* = Linear, *α_c_* = 0.03, and *α_p_* =0.03).

**Fig. 5.**
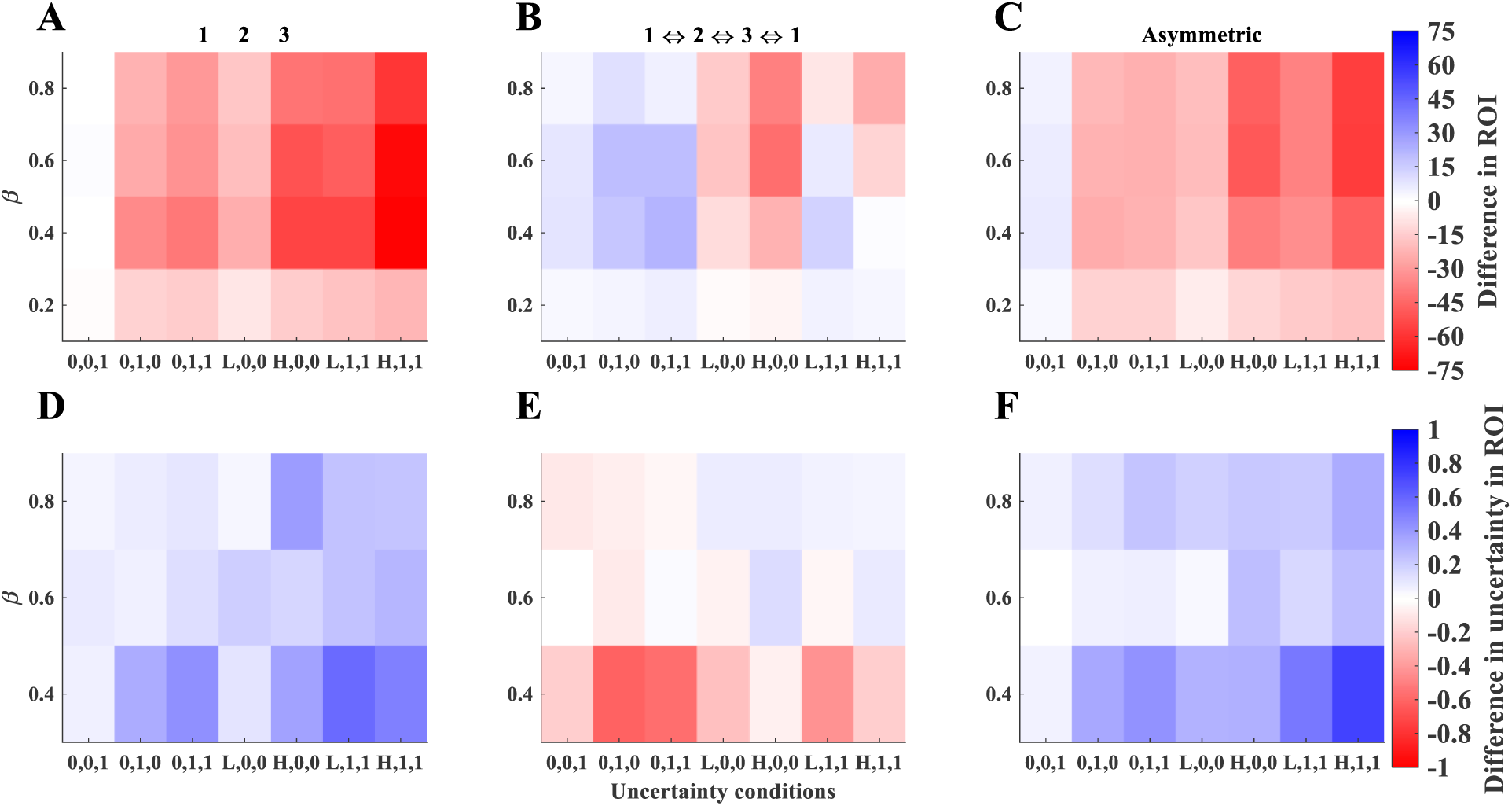
Effects of uncertainty in abundance and connectivity on ROI (A-C) and its error (D-F). X-axes define the uncertainty conditions (0,0,1 = *U_Xj→k_*; 0,1,0 = *U_X_*; 0,1,1 = *U_Xj→k_* & *U_X_*; L,0,0 = low U_a_; H,0,0 = high U_a_; L,1,1 = low U_a_ & *U_Xj→k_* & *U_X_*; H,1,1 = high U_a_ & *U_Xj→k_* & *U_X_*). A-C shows the difference in mean ROI (predictions from model in Appendix 1: Table S1) for each set of uncertainty conditions as compared with no uncertainty, whereas D-F show the difference in coefficient of variation (CV of the raw data) between conditions with uncertainty versus those without. Note, when β = 0.2 the CV’s were generally very high and stochastic even with no uncertainty in connectivity and abundance so these were omitted. True connectivity structures that were evaluated are indicated at the top of the plots; ‘asymmetric’ refers to the average of all 4 connectivity structures that were not symmetric.

In general, ROI tended to be higher when the damage function was exponential but the magnitude of effect depended on *β* (Appendix 1: Fig. S4, S5) and connectivity structure (Fig. 2B). While absolute ROI was affected by the damage function, prioritization of the source (either sequential or partial) remained optimal across all three damage functions (Fig. 3C-F) for the full range of *β* (data not shown). However, the damage function did affect the optimal strategy when connectivity was symmetric (Fig. 3A-B).

**Fig. 3.**
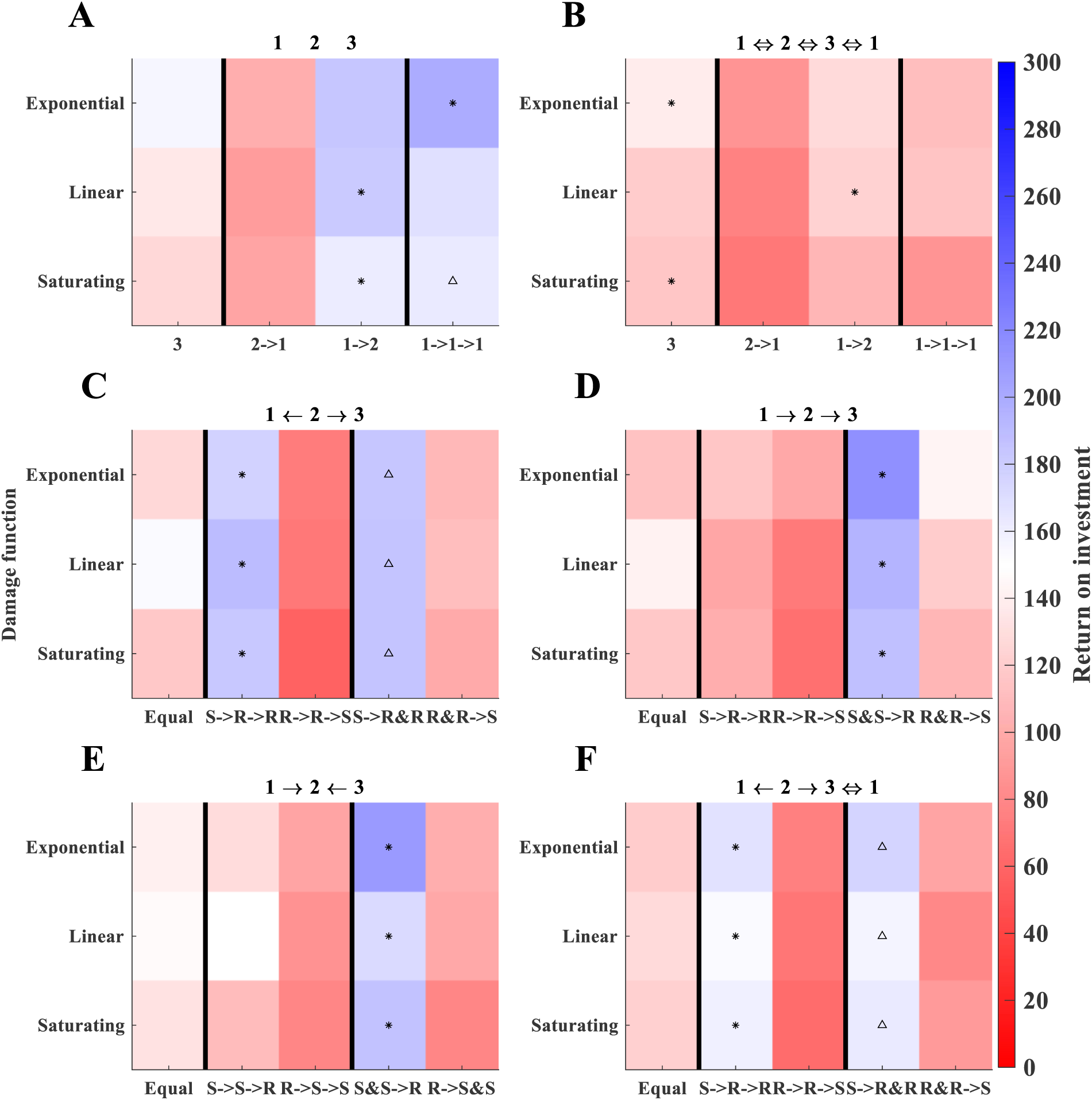
Effects of damage function on ROI and optimal strategy. Colors represent mean ROI of 100 replicate simulations. Each plot shows results for a different fixed connectivity structure. X-axes for symmetric connectivity structures (A-B) indicate the number of subpopulations that are prioritized simultaneously (e.g., 3 means that all subpopulations are equally prioritized, whereas 2→1 means that two subpopulations were initially prioritized simultaneously followed by the third). X-axes of the asymmetric connectivity structures (C-F) indicate the order of source/recipient prioritization, where ‘Equal’ refers to distributing all resources equally, & indicates that two subpopulations are being prioritized simultaneously, and → indicates sequential prioritization. Y-axes indicate the shape of the damage function. Fixed conditions were: β = 0.6, *α_c_* = 0.03 and *α_p_* =0.03, Dispersal probability = 0.1, Abundance uncertainty: none, Connectivity uncertainty: none. Stars indicate the ‘optimal’ strategy; triangles are not significantly different from the stars.

### Connectivity and dispersal probability

For asymmetric connectivity, ROI was always higher when sources were prioritized, regardless of dispersal probability values (Fig. 4). However, when subpopulations had full symmetric connectivity, equal resource allocation was best at high levels of dispersal probability, while unequal resource allocation was better at lower values of dispersal probability (Fig. 4). When there were multiple source populations, it was optimal to prioritize both sources at the same time rather than to allot resources to each sequentially before allotting resources to the recipient (Fig. 4D-E). Although the optimal strategies tended to have higher uncertainty, the absolute range of uncertainty in ROI was low relative to the large differences in mean ROI showing that the best strategies were optimal both in terms of mean and uncertainty in ROI (Fig. 4, Appendix 1: Fig. S7).

**Fig. 4.**
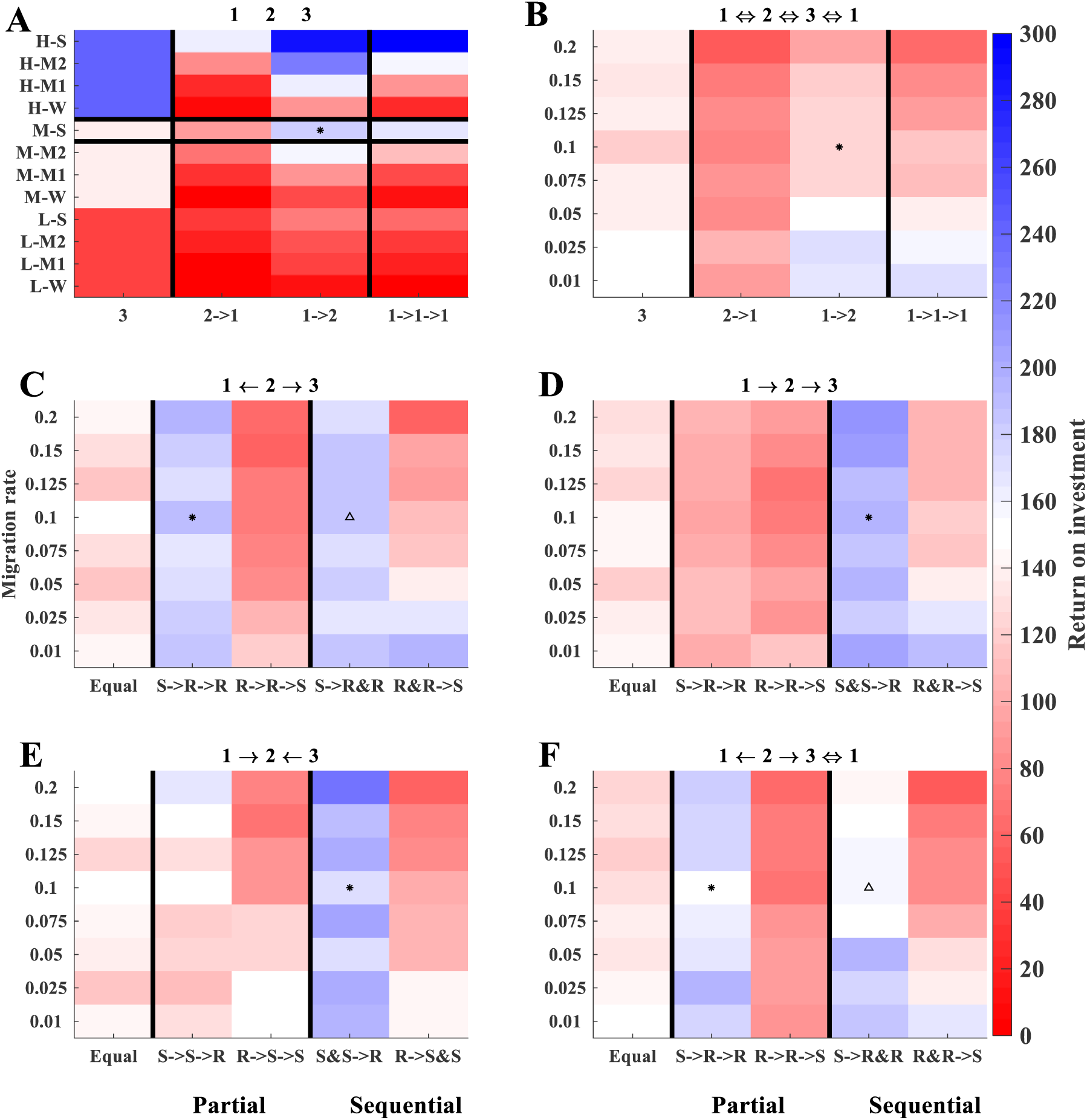
Effects of prioritization strategies on ROI (%). Colors represent mean ROI of 100 replicate simulations. Each plot shows results for a different fixed connectivity structure. X-axes for symmetric connectivity structures (A-B) indicate the number of subpopulations that are prioritized simultaneously (e.g., 3 means that all subpopulations are equally prioritized, whereas 2→1 means that two subpopulations were initially prioritized simultaneously followed by the third). X-axes of the asymmetric connectivity structures (C-F) indicate the order of source/recipient prioritization, where ‘Equal’ refers to distributing all resources equally, & indicates that two subpopulations are being prioritized simultaneously, and → indicates sequential prioritization. Y-axes are population-level annual dispersal probabilities (B-F), or levels of the α_c_ and α_p_ parameters (A, where H-, M-, L-correspond to high (0.07), medium (0.03), and low (0.013) values of capture success (α_c_) as shown in Appendix 1: Fig. S1, and S, M1, M2, W correspond to strong (0.03), moderate 2 (0.01), moderate 1 (0.005), and weak (0.0025) values of prioritization strength (α_p_) as shown in Appendix 1: Fig. S2). The horizontal black lines in A show results for the values of *α_c_* = 0.03 and *α_p_* =0.03 that are used in plots B-F (note condition A has no potential for dispersal). Other fixed conditions were: β = 0.6, *f_d_* = Linear, Abundance uncertainty: none, Connectivity uncertainty: none. Stars indicate the ‘optimal’ strategy under these conditions and the average dispersal probability when there is no uncertainty; triangles are not significantly different from the stars.

### Uncertainty

Uncertainty in abundance and connectivity structure led to lower ROI (Fig. 5A-C) with higher uncertainty in ROI (Fig. 5D-F), but the magnitude of effects on mean ROI depended on control effort (Fig. 5, Appendix 1: Fig. S8, Appendix 1: Table S1 rows 34-36). The effects of uncertainty in abundance and connectivity structure tended to be strongest at moderate values of control effort (β=0.4 or 0.6; Fig. 5, Appendix 1: Fig. S9). When all types of observational uncertainty were present and β = 0.6, ROI was 57% and 73% lower on average in cases where the true connectivity was none or asymmetric. Uncertainty in connectivity rate was less important than uncertainty in connectivity structure and abundance (Fig. 5, Appendix 1: Fig. S8, S9). The negative effects of uncertainty in abundance were not significantly different across true connectivity structures (Appendix 1: Table S1 rows 19-23), while the magnitude of effects of uncertainty in connectivity structure and rate on ROI depended significantly on the true connectivity structure (Fig. 5, Appendix 1: Table S1 rows 24-33). There was a significantly negative interaction between uncertainty in connectivity rate and structure (Appendix 1: Table S1 row 18), but no significant interaction between uncertainty in abundance and connectivity structure or rate (Appendix 1: Table S1 rows 16-17).

## DISCUSSION

Ecological and bioeconomic theory suggests that spatial prioritization of resources is important for efficient control of invasive species or disease (Hanski and Ovaskainen 2000, Chades et al. 2011, Hastings 2014, Baker 2017). However, applying spatial prioritization plans in practice requires data that describe the connectivity between subpopulations in space. As quantifying demographic connectivity directly can be time-consuming and expensive, one way to infer movement between subpopulations is to collect genetic samples from the subpopulations and analyze the data using population genetic approaches (Lowe and Allendorf 2010). Here we showed that uncertainty in directionality of dispersal can substantially reduce ROI and increase its uncertainty. Our results highlight an important challenge for practitioners and geneticists because current methodologies for estimating connectivity using genetics can be highly uncertain or biased unless true demographic processes meet specific conditions (Meirmans 2014, Samarasin et al. 2017), and connectivity structure can be dynamic. The silver lining is that uncertainty in dispersal rates seemed less important than uncertainty in dispersal direction, and genetic methods typically have less uncertainty in estimating dispersal direction than in estimating dispersal rates (Samarasin et al. 2017).

In practical applications, it is important to understand how budget constraints affect the optimal control strategy in order to use the available resources efficiently (Epanchin-Niell and Hastings 2010). We found that it was better to prioritize source populations, similar to Baker et al. (2017), across a range of monthly budgets when connectivity was asymmetric. However, the optimal resource allocation strategy was affected by budget under no connectivity due to the non-linear relationship between capture-success and abundance. In these cases, some form of prioritization was always advantageous but whether the strategy should be partial or sequential depended on budget. Our finding that the budget can affect not just the amount of damage prevented but also the strategy that would lead to the highest ROI and its uncertainty is consistent with previous work based on empirical data from willow (*Salix cinerea*) invasion management in the Australian Alps (Giljohann et al. 2011). For example, Giljohann et al. (2011) found that the budget affected which sites should be included in the control plan as well as time investment in each site. In general, under low budgets and high connectivity (symmetric or asymmetric), prioritization had little advantage over equal allocation of resources because elimination probability was low for all prioritization schemes under these conditions. Thus, ultimately the optimal resource allocation strategy depends most strongly on whether there are enough resources to rapidly reduce each subpopulation and dispersal probabilities when prioritization is applied (i.e., removal rates that are well beyond birth and immigration rates).

Another important factor determining the optimal resource allocation strategy was the damage function. Previous work has demonstrated that ROI from control can vary dramatically depending on the damage function and the relative difference of damage costs to control costs (Yokomizo et al. 2009, Davis et al. 2018a). Specifically, when the damage function is saturating and damage costs are low to moderate, there can be little to gain by investing in control (Yokomizo et al. 2009, Davis et al. 2018a). Similarly, we also found that ROI for the optimal strategy was generally lower when the damage function was saturating or linear as compared to exponential. As with the effects of budget, we found that the damage function only affected the optimal resource allocation strategy when connectivity was symmetric, although it did affect absolute ROI under all connectivity structures. Thus, the damage function plays an important role in determining overall ROI, and the optimal strategy depending on the true connectivity structure. Future work should evaluate the ROI of allocating resources into monitoring to decrease uncertainty in the damage function, e.g., see (Moore and Runge 2012) for a similar concept.

A third important factor affecting the optimal strategy was observational uncertainty in abundance and connectivity. Our model assumed that the manager allocated resources using a pre-determined prioritization scheme based on knowledge of connectivity structure and using monthly estimates of subpopulation abundances. Thus, when uncertainty in subpopulation abundance was high, subpopulations may have received more or fewer resources than if abundance had been observed more accurately. This allocation error resulted in decreased and more uncertain ROI, similar to (Bonneau et al. 2019, Regan et al. 2019), and occurred regardless of the true connectivity structure. Uncertainty in connectivity did not explain as much of the variation in ROI for different resource allocation strategies as compared to uncertainty in abundance. However, when uncertainty in connectivity structure was present, higher control intensities were needed in order to have more certainty about ROI of the optimal strategy. This was because when the connectivity structure was uncertain, there was error in identifying the true source population(s) such that benefits from reducing dispersal probabilities were reduced. Also, when the true connectivity structure was ‘completely connected’, ROI was higher and less uncertain when there was uncertainty in connectivity structure (data not shown). This was because ROI in several other connectivity structures tended to be higher than ROI in fully connected populations across several different prioritization strategies (i.e., the fully connected case was generally a worst case scenario such that if connectivity was less than fully connected applying any strategy would still give a higher ROI). In contrast, uncertainty in connectivity structure generally decreased expected ROI and increased its uncertainty for all other types of true connectivity structures. Thus, uncertainties from both abundance and demographic connectivity are important for determining optimal resource allocation strategies. Incorporating both sources of demographic uncertainty, in a manner that appropriately links and propagates their uncertainties will be important for determining optimal resource allocation strategies in practice.

While our model accounted for several realistic non-linear relationships of management costs and density (e.g., similar to (Baker 2017)), we did not consider density-dependent dispersal between subpopulations, nor dispersal outside the original area occupied by the metapopulation. Instead, we modeled dispersal as a fixed proportion of the source population and without regard to recipient population density. While density-independent dispersal could be unrealistic in some contexts (Olsson et al. 2006, Strevens and Bonsall 2011, Tabak et al. 2017, De Bona and Bruneaux 2019), density-independent connectivity could occur in some situations where anthropogenic movement shapes the connectivity structure (Hernandez et al. 2018). Regardless, we expect that incorporating density-dependent dispersal may have only a minor impact on the relative importance of factors affecting the optimal resource allocation strategy because we found that the details of connectivity structure were less important than the overall control effort and uncertainty in abundance. Although patterns of dispersal were density-independent, we assumed that birth rates in local populations were density-dependent. Limiting subpopulations by density-dependent births could affect the optimal prioritization strategy because uncontrolled populations would be allowed to expand further, causing more damage, and resource allocation decisions and damage functions are dependent on densities. Thus exponential growth could offset the benefits of spatial prioritization (depending on spatial expansion rates and damage costs), e.g., (Bonneau et al. 2017). Exponential growth in local populations could be realistic in the case of wild pigs in the U.S.A., where uncontrolled populations continue to expand more than controlled populations (Pepin et al. 2019), i.e., they are not limited by favorable habitat. An understanding of density dependence in demographic processes will be important for applying connectivity-based management on real landscapes.

We tested the concept that managers could use a one-time estimate of demographic connectivity that was obtained at the outset of a management program to identify an optimal resource allocation scheme. We then tested how uncertainty in that one-time estimate would affect robustness of the optimal resource allocation strategy that is applied. We assumed that for a given scenario, connectivity structure and rate were fixed. As connectivity is likely to change due to management or environmental factors, developing bioeconomic frameworks that account for these effects, and how additional surveillance and learning could optimize connectivity-based management plans, will be important for realizing the full benefits of connectivity-based management in practice.

Our results highlighted the importance of accurate monitoring data for planning optimal resource allocation strategies – uncertain knowledge of demographic processes and economic relationships can result in choosing resource allocation strategies with lower ROIs. However, we did not consider the costs of different types of monitoring data (e.g., abundance, connectivity, damage costs). An important future direction is to determine how much investment in monitoring demographic and economic processes would be needed (if any) to maximize ROI by increasing the confidence in choosing the optimal connectivity-based control strategy (Epanchin-Niell et al. 2012, Moore and Runge 2012, Rout et al. 2014, Baker and Bode 2016, Gormley et al. 2016). Our results suggest that robust estimates of connectivity will be especially important when connectivity is asymmetric, and that reduced uncertainty in economic data is most important when connectivity is symmetric. Future work that considers the monitoring costs for reducing uncertainties in the data used to guide management decisions is important for determining how resources can optimally be apportioned to control and monitoring in realistic contexts (i.e., where uncertainty in demographic dynamics and damage functions exist).

## Supporting information

Code for model

## AUTHOR CONTRIBUTIONS

KMP coded the model, ran simulations, analyzed output, and wrote the first draft. All other authors contributed to concept development and manuscript editing.

## ACKNOWLEDGEMENTS

We thank Gabriel Gellner, David Wolfson, and Michael Tabak for helpful discussion during the development of this work, as well as two anonymous reviewers whose comments greatly improved the manuscript. KMP, AJD, TJS, CS, and SM were funded by the United States Department of Agriculture, Animal and Plant Health Inspection Service’s National Feral Swine Damage Management Program.

## Appendix 1

**Figure S1.**
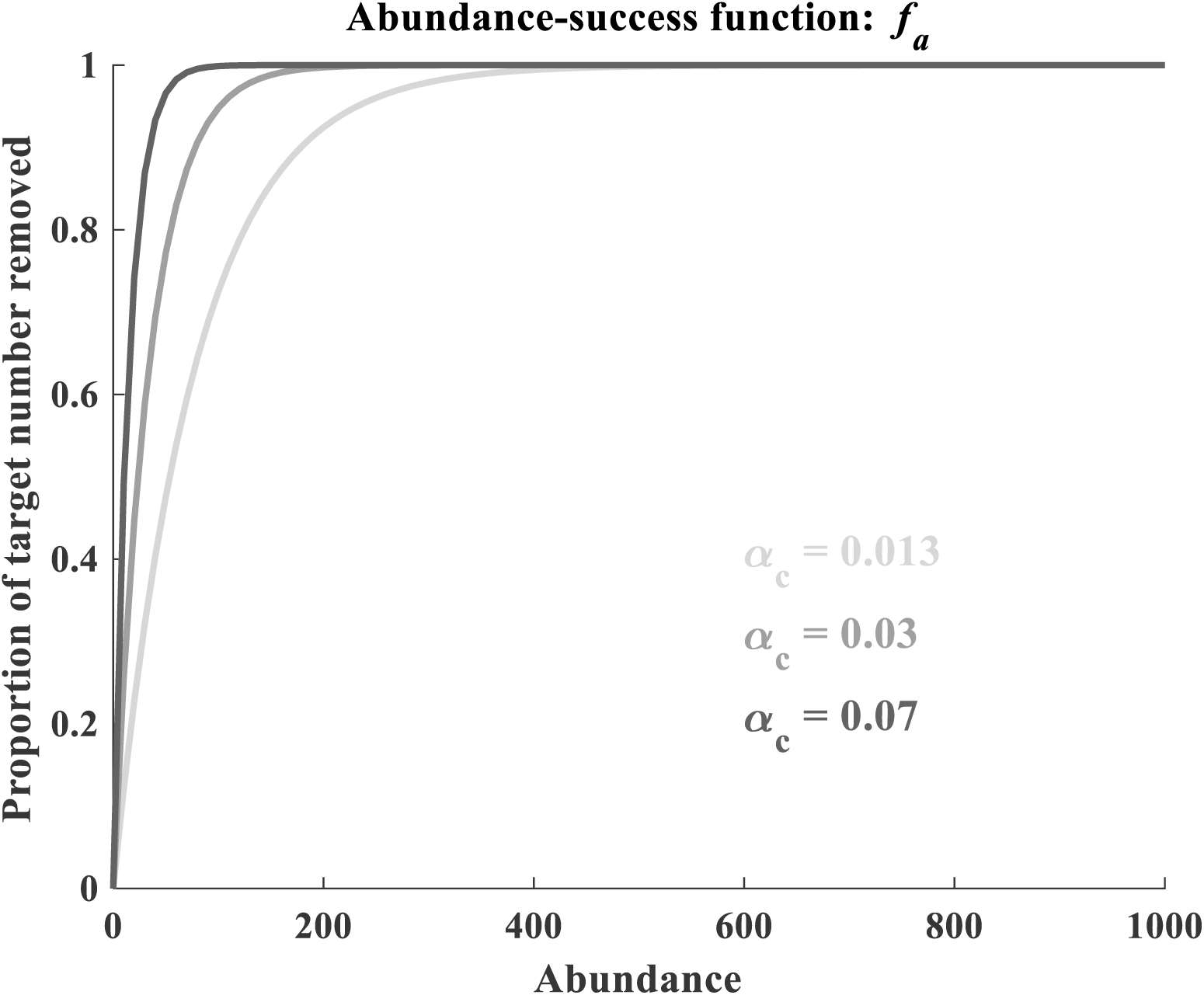
The abundance-success function, where N is abundance, *t* is month, *k* indicates the subpopulation, and α_c_ is a scaling parameter. 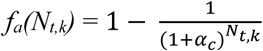

**Figure S2.**
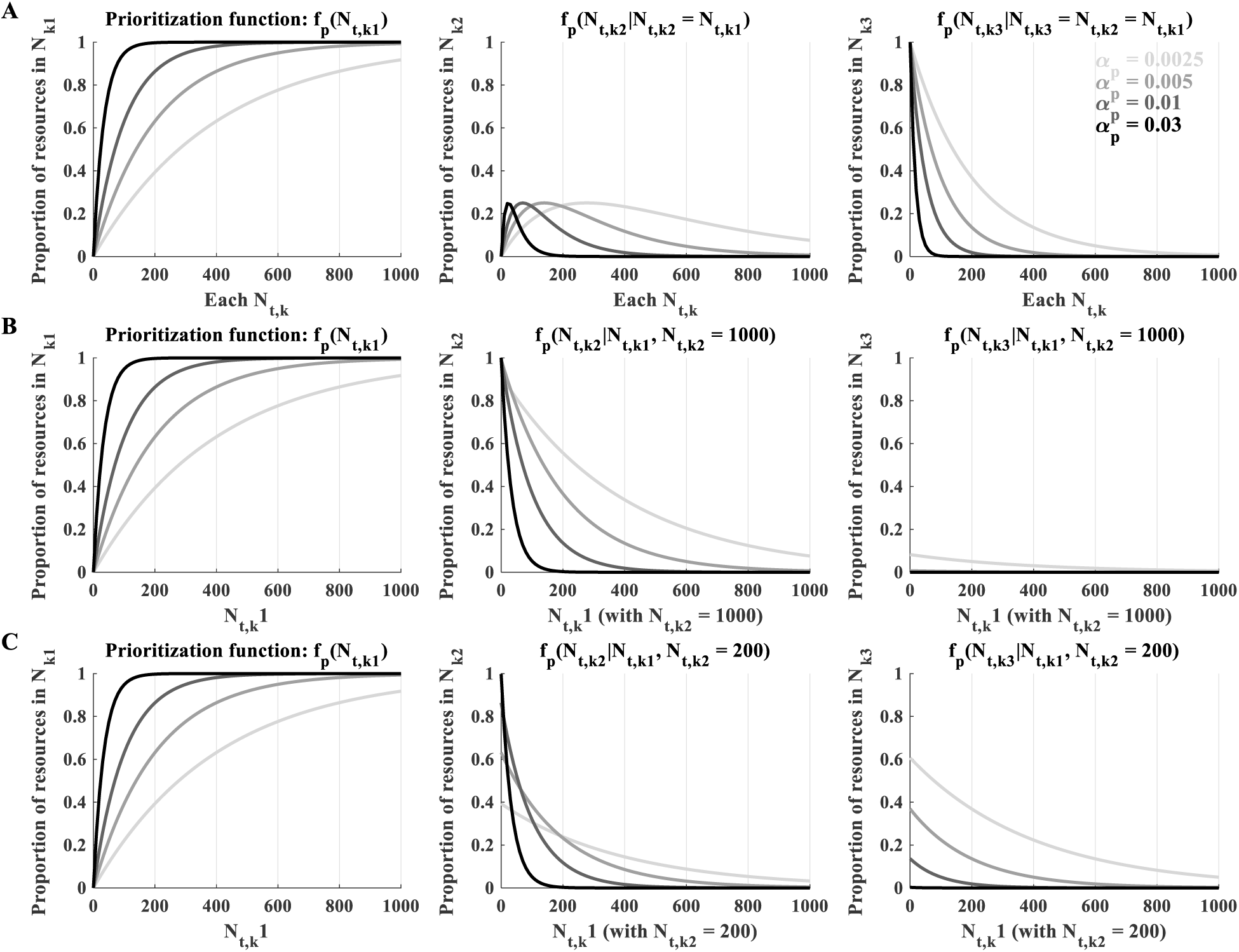
Prioritization functions given different examples of abundance in the first and second most prioritized subpopulations. Row A shows the proportion of abundances allotted to each subpopulation under the condition where all subpopulations are at the same abundance (this should only occur at the outset when all subpopulations = 1000; but the plots serve as an example for visualizing the prioritization functions). Different lines within the plots show how the function differs depending on different values of the scaling parameter (*α_p_* – values in upper right plot). Rows B and C are similar to row A but show *f_p_* for each subpopulation given N_t,k1_ on the X-axis when N_t,k2_ = 1000 (B) or 200 (C). The last priority subpopulation always receives the rest of the resources and thus depends exclusively on N_t,k1_ and N_t,k2_, rather than its own abundance.

**Figure S3.**
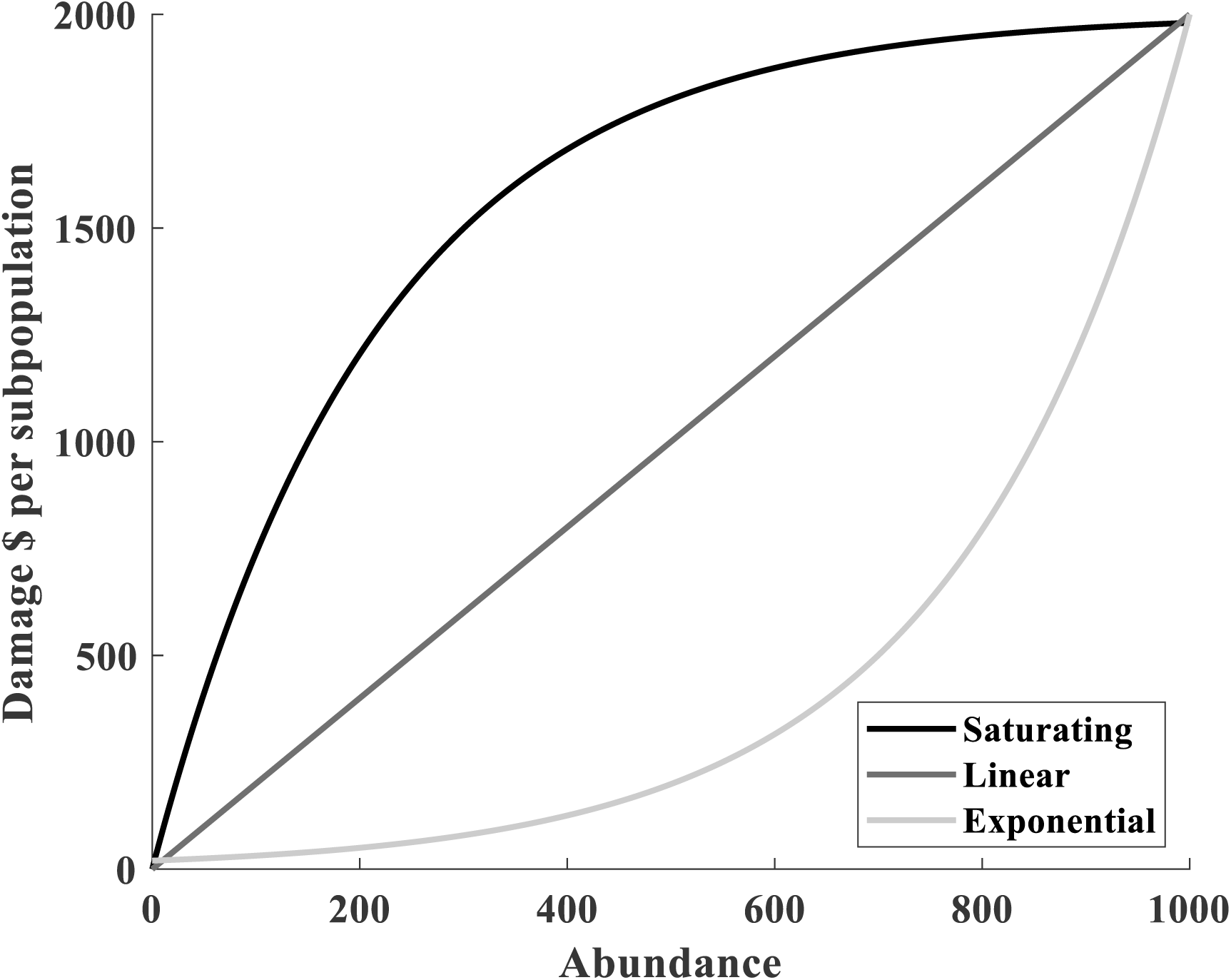
Shape of the three damage functions – saturating (Type II), linear (Type I), and exponential (Type IV) from Eq.s 11-13 in the main text..

**Figure S4.**
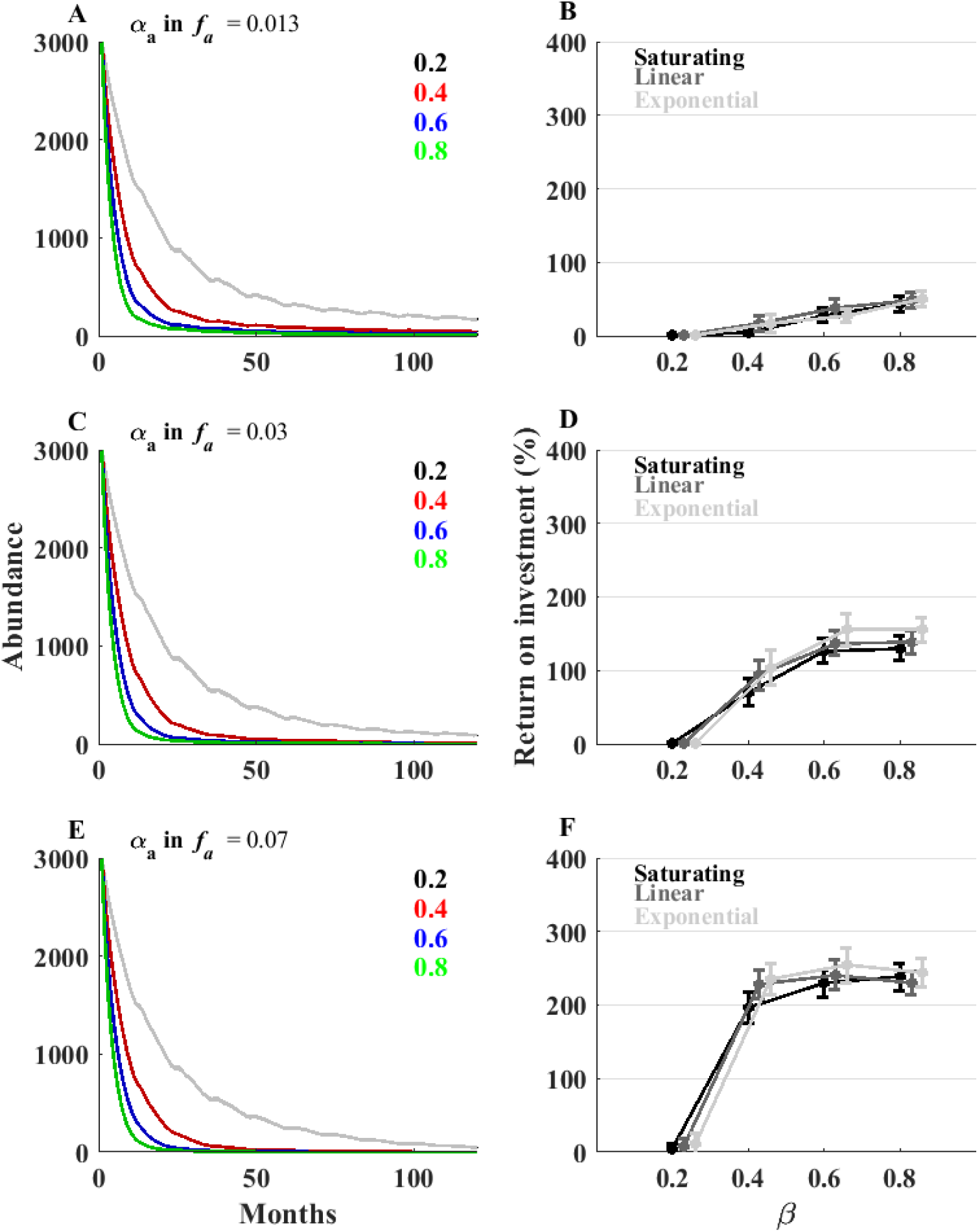
Behavior of the model with no connectivity or prioritization. A. Effects of control intensity on abundance over 10 years. Control intensity is the maximum proportion of the population that can theoretically be removed per month. Actual numbers depend on the abundance, where removal becomes more difficult as abundance decreases. Lines are means of 100 replicate simulations with 95% confidence intervals (indicated by shading around the means – 0.2 (grey), 0.4 (red), 0.6 (blue), 0.8 (green); not easily visible because CI’s are tight). B. Effects of control intensity and the relationship of population density and damage costs on ROI over 10 years. ROI is shown as a function of control intensity for 3 assumptions about the shape of the relationship between population density and damage costs: saturating, linear, exponential. The three sets of paired plots show results for the three different values of *α_a_* (indicated at the top of A figures). Fixed conditions were as in Table 1 main text. We also fixed the following variable input parameters: X = Connectivity structure 1 (unconnected), *X_j→k_* = 0, *f_p_* = no prioritization, *f_d_* = Linear, ζ=0.5, Abundance uncertainty: none, Connectivity uncertainty: none.

**Figure S5.**
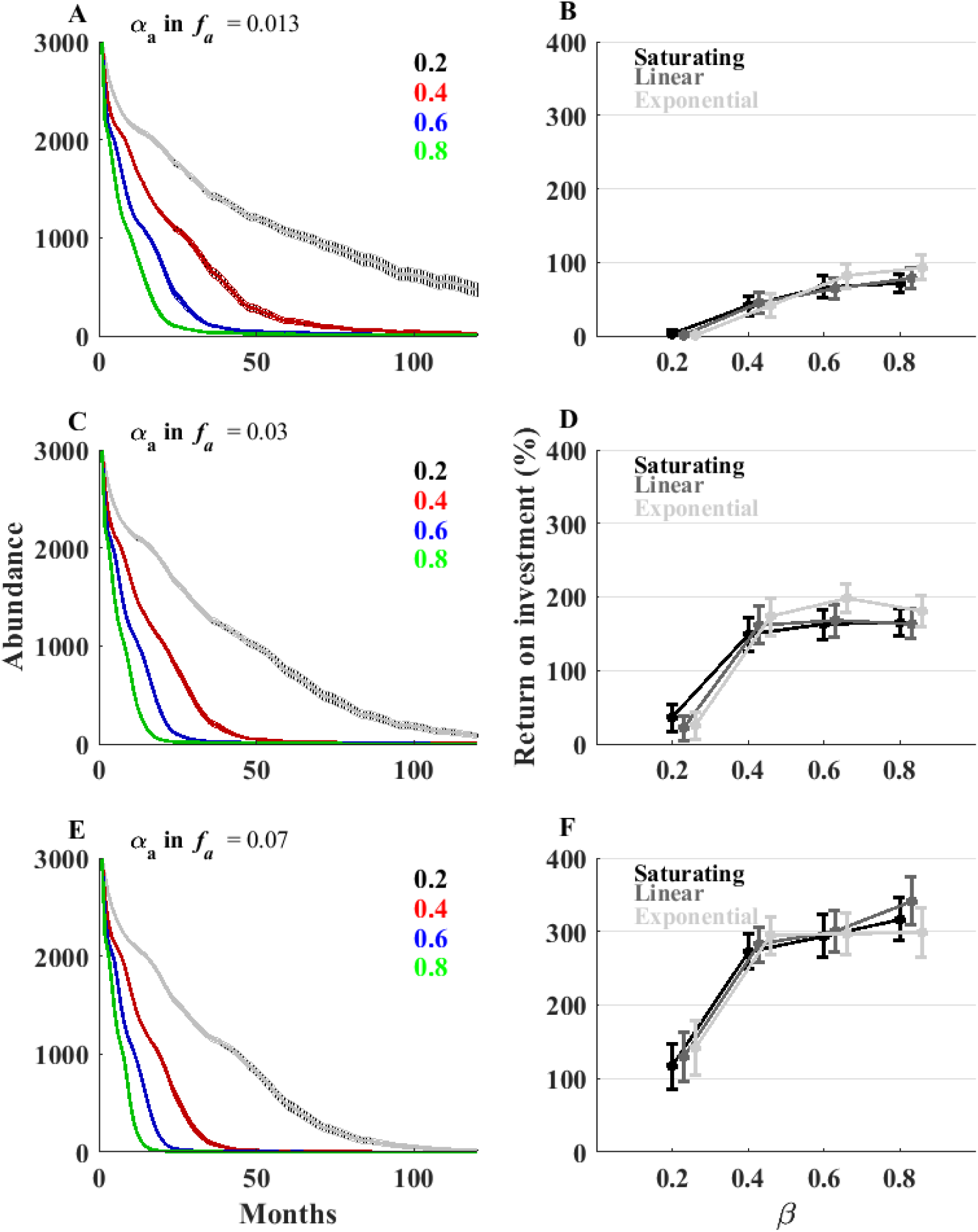
Same as Fig. S4 but with sequential prioritization of subpopulations.

**Fig S6.**
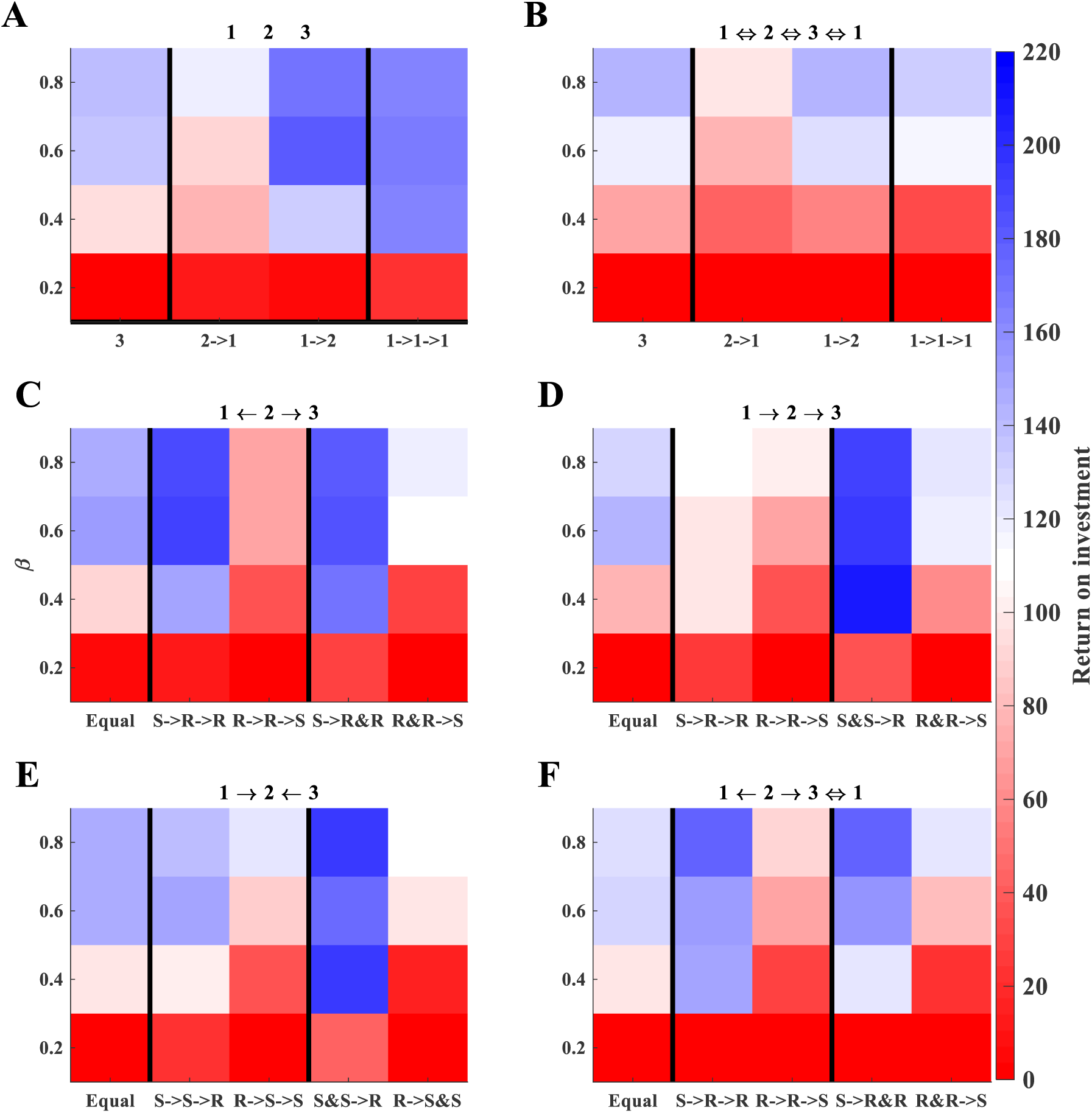
Same as Fig. 3 in the main text except Y-axes are control effort rather than dispersal probability. Other fixed conditions were: dispersal probability = 0.1, *f_d_* = Linear, *α_c_* = 0.03, and *α_p_* = 0.03, Abundance uncertainty: none, Connectivity uncertainty: none.

**Fig. S7.**
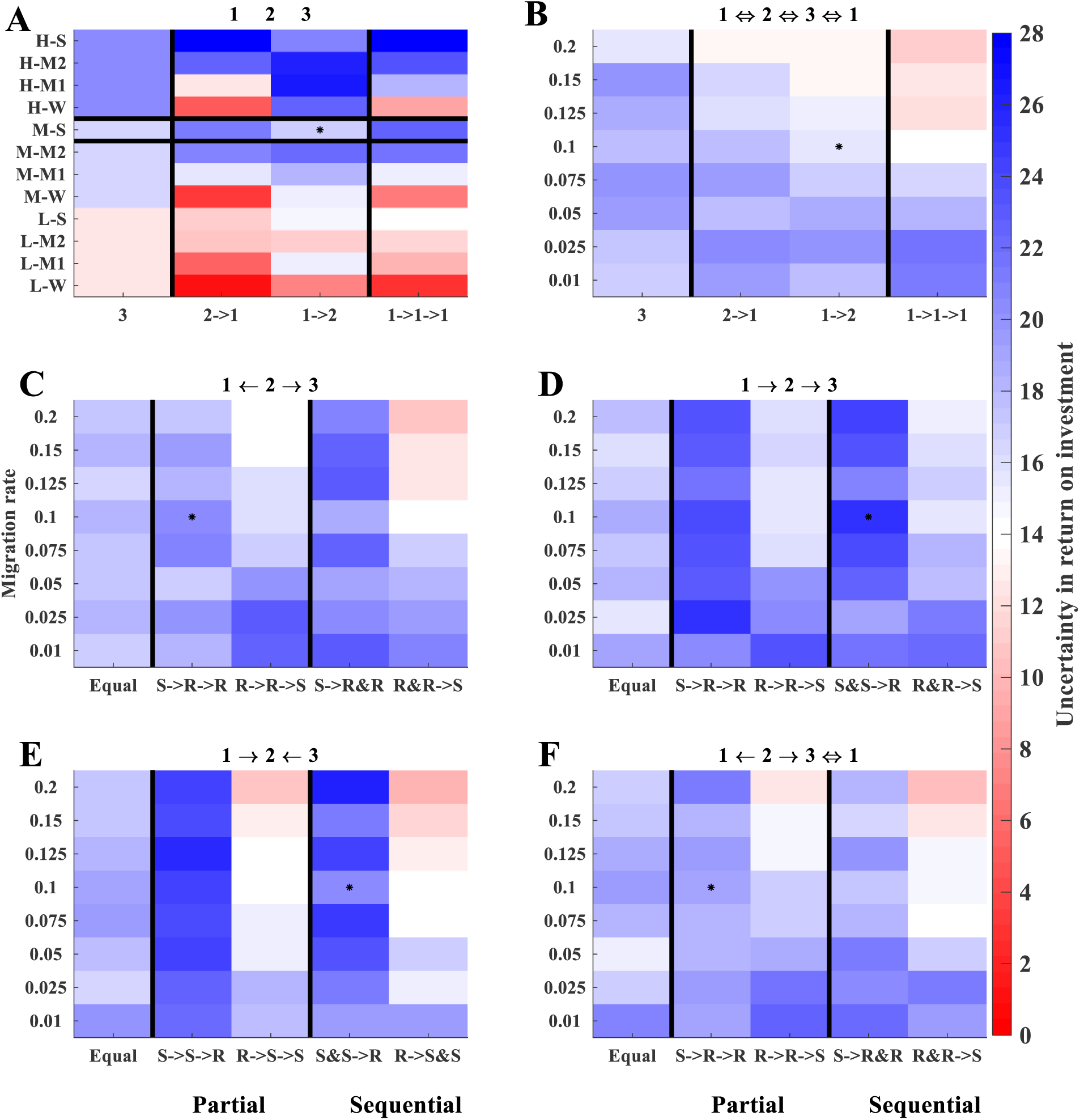
Effects of prioritization strategies on uncertainty in ROI (%). Colors represent 95% confidence intervals of mean ROI from 100 replicate simulations. Each plot shows results for a different fixed connectivity structure. X-axes for symmetric connectivity structures (A-B) indicate the number of subpopulations that are prioritized simultaneously (e.g., 3 means that all subpopulations are equally prioritized, whereas 2→1 means that two subpopulations were initially prioritized simultaneously followed by the third). X-axes of the asymmetric connectivity structures (C-F) indicate the order of source/recipient prioritization, where ‘Equal’ refers to distributing all resources equally, & indicates that two subpopulations are being prioritized simultaneously, and → indicates sequential prioritization. Y-axes are population-level annual dispersal probabilities (B-F), or levels of the α_c_ and α_p_ parameters (A, where H-, M-, L-correspond to high (0.07), medium (0.03), and low (0.013) values of capture success (α_c_) as shown in Fig. S1, and S, M1, M2, W correspond to strong (0.03), moderate 2 (0.01), moderate 1 (0.005), and weak (0.0025) values of prioritization strength (α_p_) as shown in Fig. S2). The horizontal black lines in A show results for the values of *α_a_* = 0.03 and *α_p_* =0.03 that are used in plots B-F (note condition A has no potential for dispersal). Other fixed conditions were: β = 0.6, *f_d_* = Linear, Abundance uncertainty: none, Connectivity uncertainty: none. Stars indicate the ‘optimal’ strategy under these conditions when there is no uncertainty in abundance and connectivity (i.e., shown in Fig. 3 main text).

**Fig. S8.**
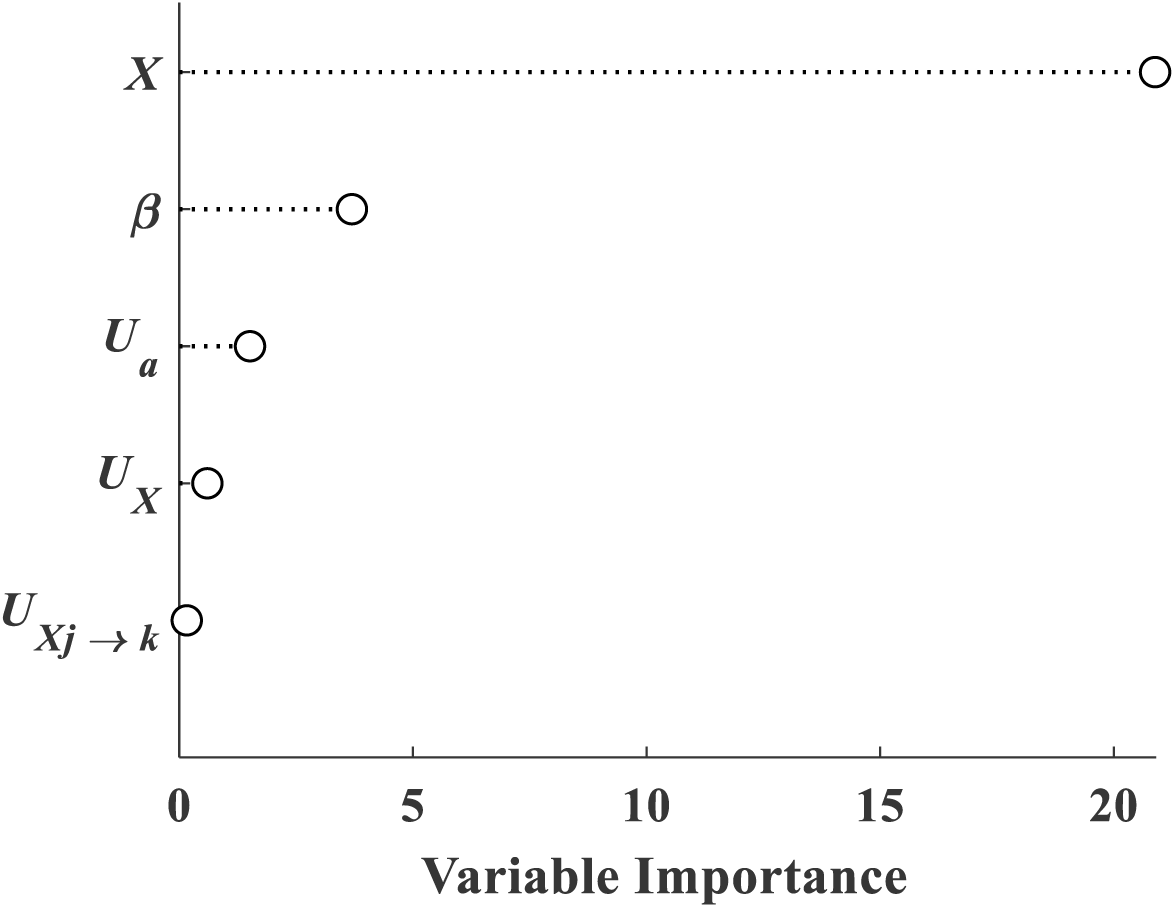
Variable importance for effects of uncertainty. We simulated effects of uncertainty in abundance and connectivity for each ‘true’ connectivity structure and the 4 values of β. We simulated all possible combinations of uncertainty sources from none to all three. We used a random forest algorithm (Breiman, L. 2001. Random Forests. Machine Learning 45:5-32) to estimate variable importance implemented using the ‘fitrensemble’ and ‘predictorImportance’ functions in Matlab R2018a (MathWorks, Inc., Natick, MA). We used bootstrap aggregating for the ensemble aggregation method, with 100 learning cycles and a maximum number of 10 splits during learning. True connectivity structure = X; Uncertainty in abundance = U_a_; Uncertainty in connectivity structure = U_X_; Uncertainty in dispersal probability = U_Xj→k_.

**Table S1.**
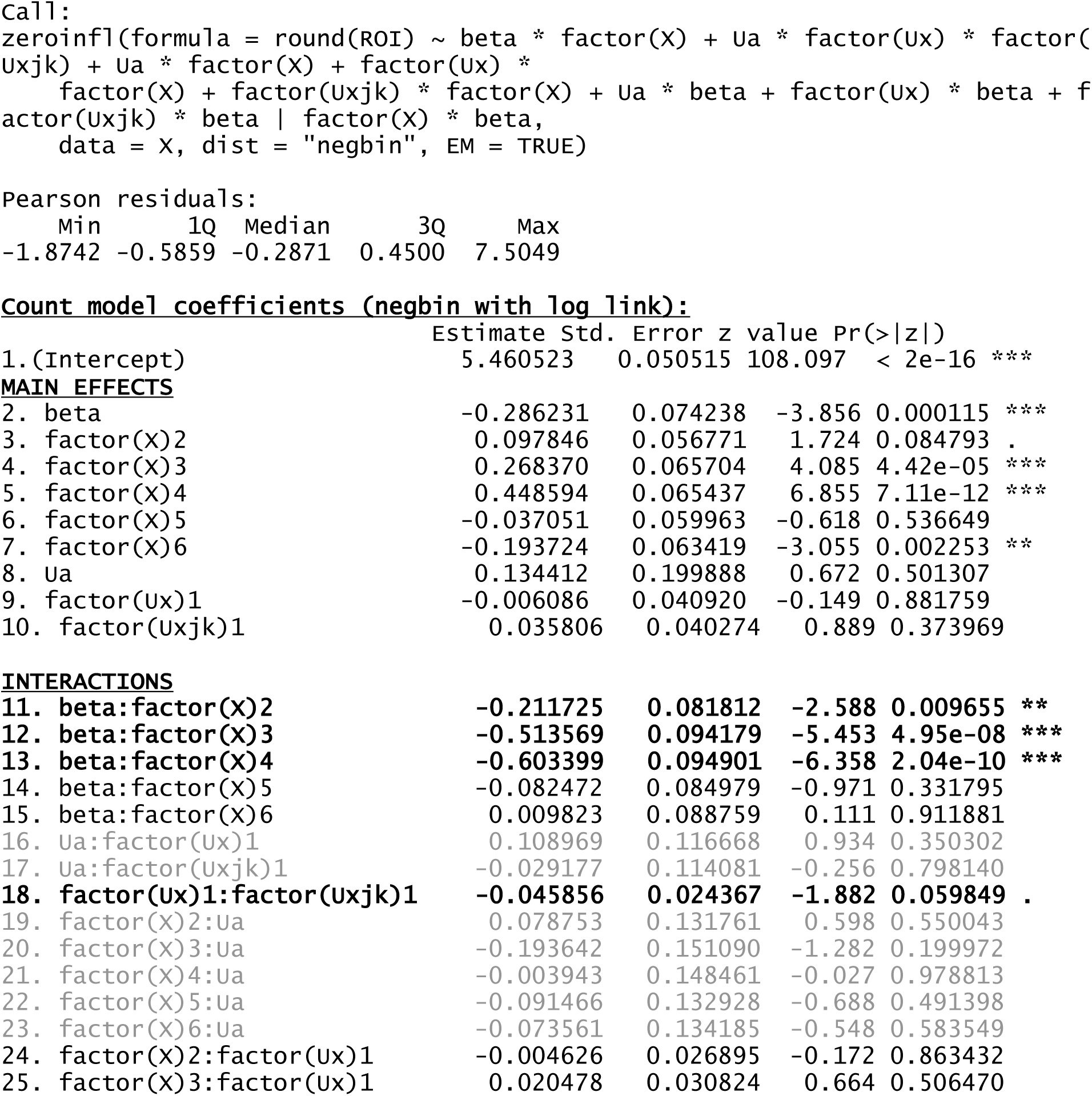
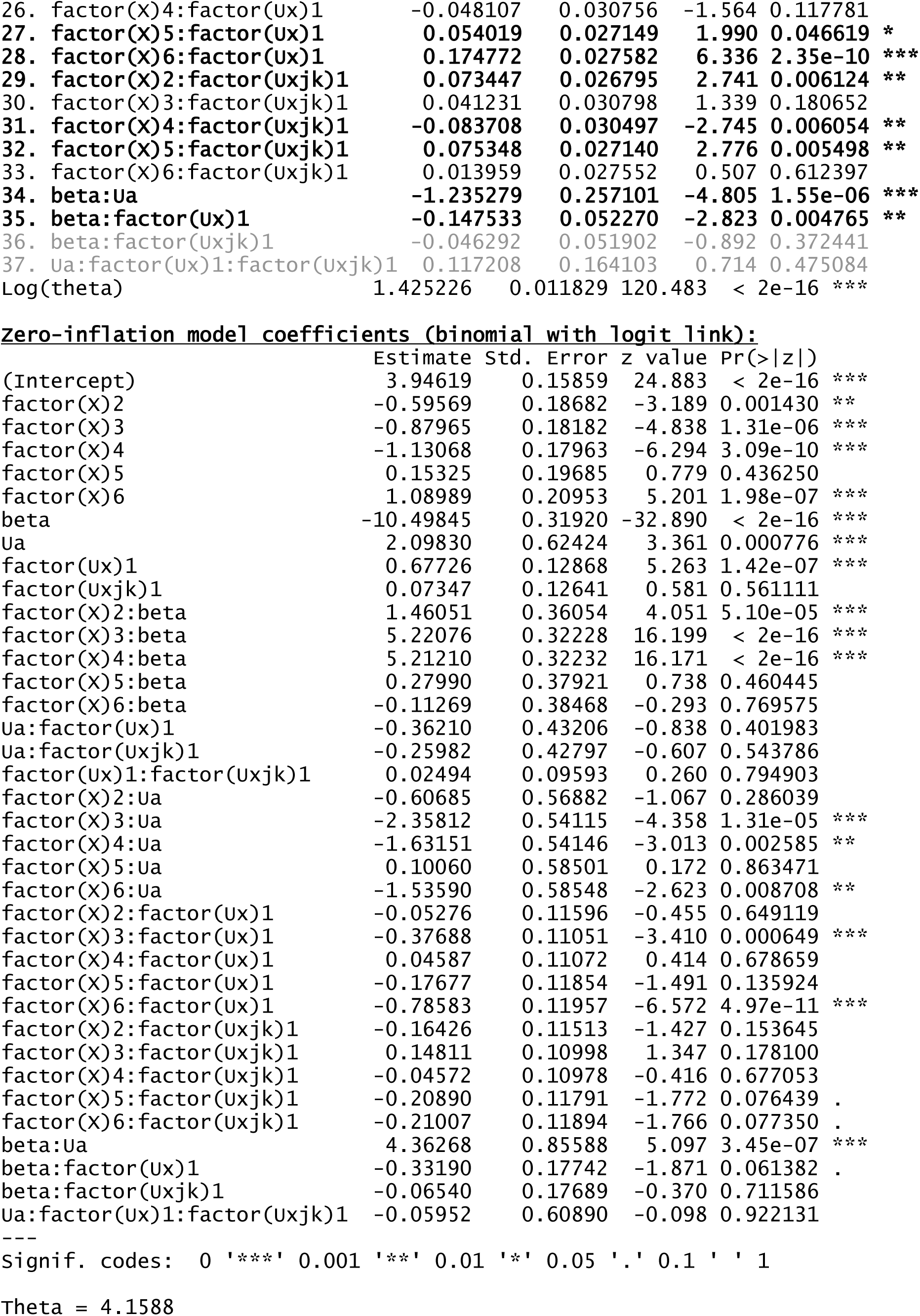
Coefficient estimates for zero-inflated negative binomial model evaluating the effects of β, connectivity structure (X), and uncertainty in abundance and connectivity on ROI. For both the negative binomial count component and the binomial component we tested the effects of interactions among sources of uncertainty, between X and β, and between each source of uncertainty (Ua = uncertainty in abundance; Ux = uncertainty in connectivity structure; Uxjk = uncertainty in dispersal probability) and each of β and X. Grey indicates interaction factors that are not significant. Bold indicates the levels of interaction factors that are significant. We used a zero-inflated negative binomial model because ROI was determined by 2 processes (i.e., a mixture): 1) ROI = 1 when elimination occurred and 1 otherwise, 2) ROI was a continuous variable > 0 when elimination occurred. Because ROI was generally much larger than 1, we took its discrete approximation by rounding to the nearest integer. We used a negative binomial model instead of a Poisson because the count data were overdispersed. R^2^ was calculated as the squared correlation between the observed and predicted data, and was 0.25. The analysis was conducted in program R using the ‘pscl’,’boot’, and ‘MASS’ packages.

Count model coefficients (negbin with log link):
|  | Estimate | Std. Error | z value | Pr(>\|z\|) |
| --- | --- | --- | --- | --- |
| 1. (Intercept) | 5.460523 | 0.050515 | 108.097 | < 2e-16 *** |
| **MAIN EFFECTS** |  |  |  |  |
| 2. beta | -0.286231 | 0.074238 | -3.856 | 0.000115 *** |
| 3. factor(X)2 | 0.097846 | 0.056771 | 1.724 | 0.084793 . |
| 4. factor(X)3 | 0.268370 | 0.065704 | 4.085 | 4.42e-05 *** |
| 5. factor(X)4 | 0.448594 | 0.065437 | 6.855 | 7.11e-12 *** |
| 6. factor(X)5 | -0.037051 | 0.059963 | -0.618 | 0.536649 |
| 7. factor(X)6 | -0.193724 | 0.063419 | -3.055 | 0.002253 ** |
| 8. Ua | 0.134412 | 0.199888 | 0.672 | 0.501307 |
| 9. factor(Ux)1 | -0.006086 | 0.040920 | -0.149 | 0.881759 |
| 10. factor(Uxjk)1 | 0.035806 | 0.040274 | 0.889 | 0.373969 |
| **INTERACTIONS** |  |  |  |  |
| 11. beta: factor(X)2 | -0.211725 | 0.081812 | -2.588 | 0.009655 ** |
| 12. beta: factor(X)3 | -0.513569 | 0.094179 | -5.453 | 4.95e-08 *** |
| 13. beta: factor(X)4 | -0.603399 | 0.094901 | -6.358 | 2.04e-10 *** |
| 14. beta: factor(X)5 | -0.082472 | 0.084979 | -0.971 | 0.331795 |
| 15. beta: factor(X)6 | 0.009823 | 0.088759 | 0.111 | 0.911881 |
| 16. Ua: factor(Ux)1 | 0.108969 | 0.116668 | 0.934 | 0.350302 |
| 17. Ua: factor(Uxjk)1 | -0.029177 | 0.114081 | -0.256 | 0.798140 |
| 18. factor(Ux)1: factor(Uxjk)1 | -0.045856 | 0.024367 | -1.882 | 0.059849 . |
| 19. factor(X)2: Ua | 0.078753 | 0.131761 | 0.598 | 0.550043 |
| 20. factor(X)3: Ua | -0.193642 | 0.151090 | -1.282 | 0.199972 |
| 21. factor(X)4: Ua | -0.003943 | 0.148461 | -0.027 | 0.978813 |
| 22. factor(X)5: Ua | -0.091466 | 0.132928 | -0.688 | 0.491398 |
| 23. factor(X)6: Ua | -0.073561 | 0.134185 | -0.548 | 0.583549 |
| 24. factor(X)2: factor(Ux)1 | -0.004626 | 0.026895 | -0.172 | 0.863432 |
| 25. factor(X)3: factor(Ux)1 | 0.020478 | 0.030824 | 0.664 | 0.506470 |

|  |  |  |  |  |  |  |
| --- | --- | --- | --- | --- | --- | --- |
| 942 | 26. factor(X)4:factor(Ux)1 | -0.048107 | 0.030756 | -1.564 | 0.117781 |  |
| 943 | 27. factor(X)5:factor(Ux)1 | 0.054019 | 0.027149 | 1.990 | 0.046619 | * |
| 944 | 28. factor(X)6:factor(Ux)1 | 0.174772 | 0.027582 | 6.336 | 2.35e-10 | *** |
| 945 | 29. factor(X)2:factor(Uxjk)1 | 0.073447 | 0.026795 | 2.741 | 0.006124 | ** |
| 946 | 30. factor(X)3:factor(Uxjk)1 | 0.041231 | 0.030798 | 1.339 | 0.180652 |  |
| 947 | 31. factor(X)4:factor(Uxjk)1 | -0.083708 | 0.030497 | -2.745 | 0.006054 | ** |
| 948 | 32. factor(X)5:factor(Uxjk)1 | 0.075348 | 0.027140 | 2.776 | 0.005498 | ** |
| 949 | 33. factor(X)6:factor(Uxjk)1 | 0.013959 | 0.027552 | 0.507 | 0.612397 |  |
| 950 | 34. beta:Ua | -1.235279 | 0.257101 | -4.805 | 1.55e-06 | *** |
| 951 | 35. beta:factor(Ux)1 | -0.147533 | 0.052270 | -2.823 | 0.004765 | ** |
| 952 | 36. beta:factor(Uxjk)1 | -0.046292 | 0.051902 | -0.892 | 0.372441 |  |
| 953 | 37. Ua:factor(Ux)1:factor(Uxjk)1 | 0.117208 | 0.164103 | 0.714 | 0.475084 |  |
| 954 | Log(theta) | 1.425226 | 0.011829 | 120.483 | < 2e-16 | *** |

**zero-inflation model coefficients (binomial with logit link):**
|  | Estimate | Std. Error | z value | Pr(>\|z\|) |  |
| --- | --- | --- | --- | --- | --- |
| 957 (Intercept) | 3.94619 | 0.15859 | 24.883 | < 2e-16 | *** |
| 958 factor(X)2 | -0.59569 | 0.18682 | -3.189 | 0.001430 | ** |
| 960 factor(X)3 | -0.87965 | 0.18182 | -4.838 | 1.31e-06 | *** |
| 961 factor(X)4 | -1.13068 | 0.17963 | -6.294 | 3.09e-10 | *** |
| 962 factor(X)5 | 0.15325 | 0.19685 | 0.779 | 0.436250 |  |
| 963 factor(X)6 | 1.08989 | 0.20953 | 5.201 | 1.98e-07 | *** |
| 964 beta | -10.49845 | 0.31920 | -32.890 | < 2e-16 | *** |
| 965 Ua | 2.09830 | 0.62424 | 3.361 | 0.000776 | *** |
| 966 factor(Ux)1 | 0.67726 | 0.12868 | 5.263 | 1.42e-07 | *** |
| 967 factor(Uxjk)1 | 0.07347 | 0.12641 | 0.581 | 0.561111 |  |
| 968 factor(X)2:beta | 1.46051 | 0.36054 | 4.051 | 5.10e-05 | *** |
| 969 factor(X)3:beta | 5.22076 | 0.32228 | 16.199 | < 2e-16 | *** |
| 970 factor(X)4:beta | 5.21210 | 0.32232 | 16.171 | < 2e-16 | *** |
| 971 factor(X)5:beta | 0.27990 | 0.37921 | 0.738 | 0.460445 |  |
| 972 factor(X)6:beta | -0.11269 | 0.38468 | -0.293 | 0.769575 |  |
| 973 Ua:factor(Ux)1 | -0.36210 | 0.43206 | -0.838 | 0.401983 |  |
| 974 Ua:factor(Uxjk)1 | -0.25982 | 0.42797 | -0.607 | 0.543786 |  |
| 975 factor(Ux)1:factor(Uxjk)1 | 0.02494 | 0.09593 | 0.260 | 0.794903 |  |
| 976 factor(X)2:Ua | -0.60685 | 0.56882 | -1.067 | 0.286039 |  |
| 977 factor(X)3:Ua | -2.35812 | 0.54115 | -4.358 | 1.31e-05 | *** |
| 978 factor(X)4:Ua | -1.63151 | 0.54146 | -3.013 | 0.002585 | ** |
| 979 factor(X)5:Ua | 0.10060 | 0.58501 | 0.172 | 0.863471 |  |
| 980 factor(X)6:Ua | -1.53590 | 0.58548 | -2.623 | 0.008708 | ** |
| 981 factor(X)2:factor(Ux)1 | -0.05276 | 0.11596 | -0.455 | 0.649119 |  |
| 982 factor(X)3:factor(Ux)1 | -0.37688 | 0.11051 | -3.410 | 0.000649 | *** |
| 983 factor(X)4:factor(Ux)1 | 0.04587 | 0.11072 | 0.414 | 0.678659 |  |
| 984 factor(X)5:factor(Ux)1 | -0.17677 | 0.11854 | -1.491 | 0.135924 |  |
| 985 factor(X)6:factor(Ux)1 | -0.78583 | 0.11957 | -6.572 | 4.97e-11 | *** |
| 986 factor(X)2:factor(Uxjk)1 | -0.16426 | 0.11513 | -1.427 | 0.153645 |  |
| 987 factor(X)3:factor(Uxjk)1 | 0.14811 | 0.10998 | 1.347 | 0.178100 |  |
| 988 factor(X)4:factor(Uxjk)1 | -0.04572 | 0.10978 | -0.416 | 0.677053 |  |
| 989 factor(X)5:factor(Uxjk)1 | -0.20890 | 0.11791 | -1.772 | 0.076439 | . |
| 990 factor(X)6:factor(Uxjk)1 | -0.21007 | 0.11894 | -1.766 | 0.077350 | . |
| 991 beta:Ua | 4.36268 | 0.85588 | 5.097 | 3.45e-07 | *** |
| 992 beta:factor(Ux)1 | -0.33190 | 0.17742 | -1.871 | 0.061382 | . |
| 993 beta:factor(Uxjk)1 | -0.06540 | 0.17689 | -0.370 | 0.711586 |  |
| 994 Ua:factor(Ux)1:factor(Uxjk)1 | -0.05952 | 0.60890 | -0.098 | 0.922131 |  |

signif. codes: 0 '\*\*\*' 0.001 '\*\*' 0.01 '\*' 0.05 '.' 0.1 ' ' 1
Theta = 4.1588

**Fig. S9.**
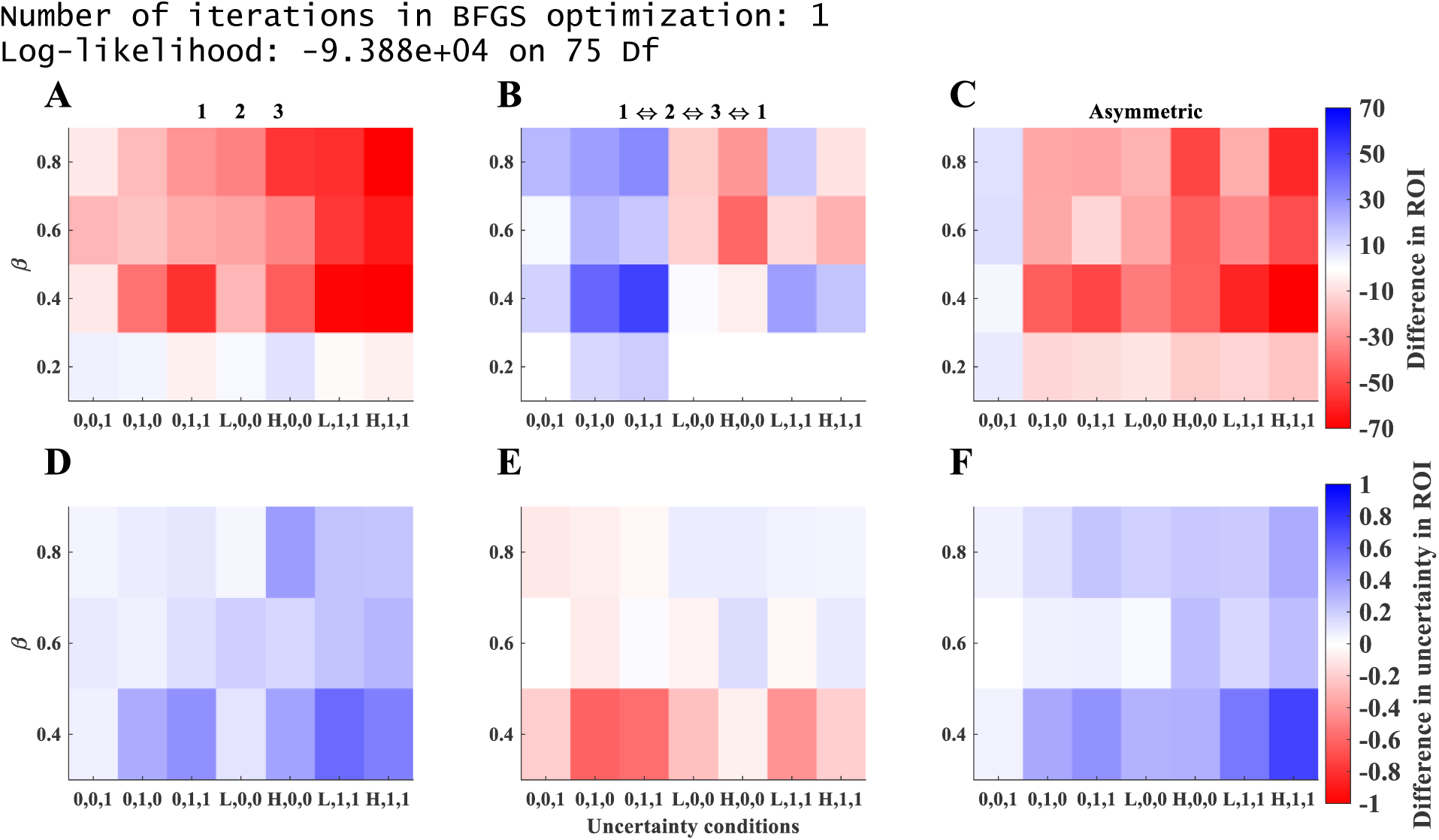
Same as Fig. 5 in the main text but for the raw data.

**Fig. S10.**
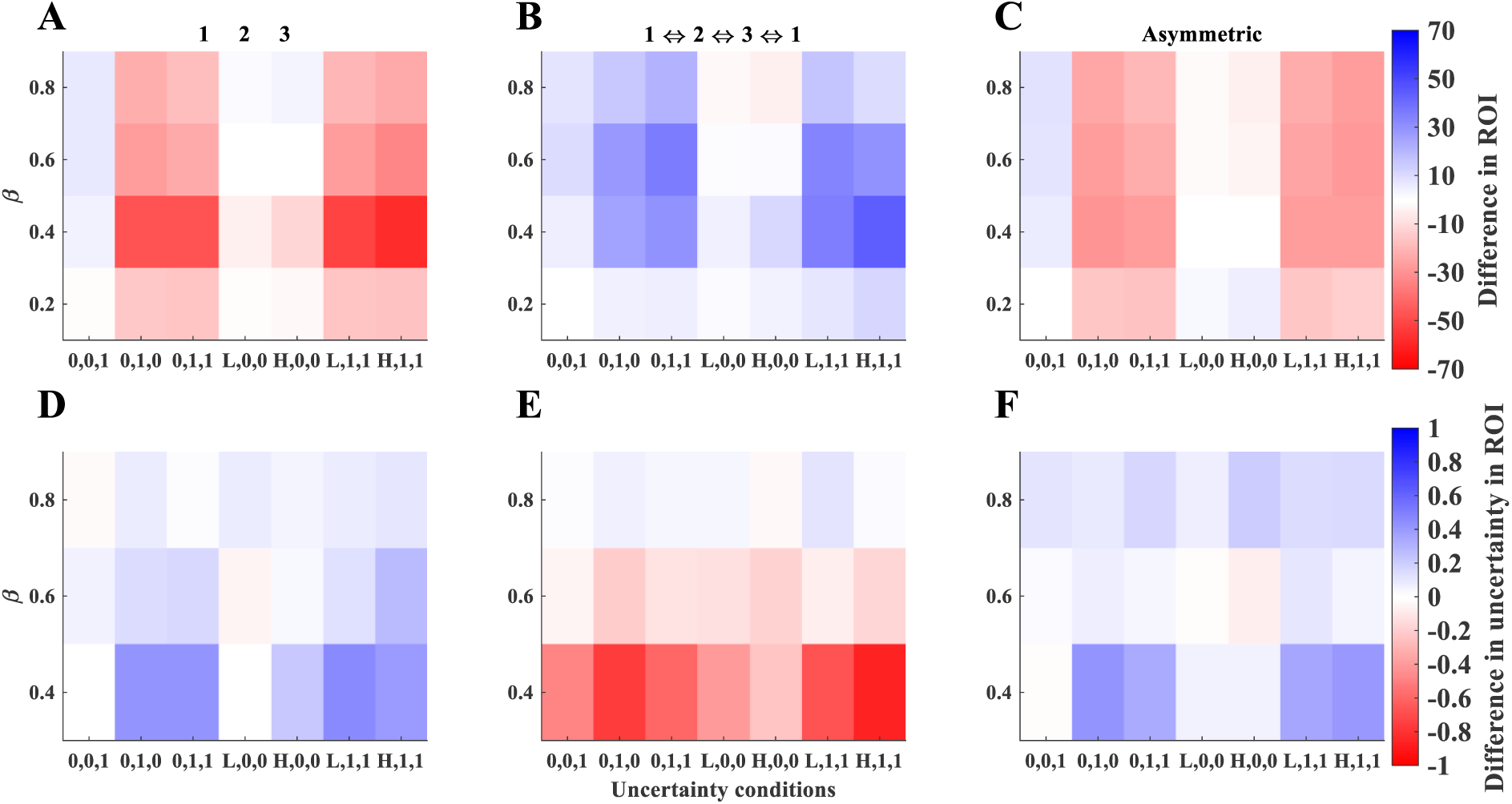
Same as Fig. 5 in the main text except here error in abundance is unbiased. The same model as in Table S1 was fit. Coefficients from the fitted model are not shown; R^2^ = 0.26.

