## Supplementary material for "Optimal spatial prioritization of control resources for elimination of invasive species under demographic uncertainty": Code for model: culling3.m

```
function effort = culling3(N,prior,beta,alpha)
% Determine how effort will be alloted across populations
% culling strategy: order of pops for culling

% Functions that describe density-dependent capture & resource allotment
% based on density
% xx = 0:0.05:1; 
% figure(101);clf; plot(xx,1-(1./(1+100).^(xx)),'b','linewidth',2);
% hold on;plot(xx,1-(1-(1./(1+100).^(1-min(1,xx)))),'r','linewidth',2); plot(xx,xx,'k','linewidth',2);
% set(gca,'box','off','fontweight','bold','fontsize',16,'fontname','times','xlim',[0 1],'ylim',[0 1]);
% ylabel('Damage costs'); xlabel('Population density');
% 
% xx = 0.01:0.05:1; yy = exp(-xx.*5)+0.2; yy = yy./(max(yy)); %1./(1-(1./(1+100).^(xx)))
% figure(101);clf; plot(xx,yy,'k','linewidth',2);
% set(gca,'box','off','fontweight','bold','fontsize',16,'fontname','times','xlim',[0 1],'ylim',[0 1]);
% ylabel('Management costs'); xlabel('Population density');

% figure(100);clf; plot(1-(1./(1+curve).^(0:100)),'k','linewidth',2);
% hold on;plot(1-(numer./(1+curve).^(0:100)),'r','linewidth',2);
% set(gca,'box','off','fontweight','bold','fontsize',12,'xlim',[0 100],'ylim',[0 1]);
% xlabel('Abundance'); ylabel('Probability of capture');
% text(103,0.9,'Proportion of resources allotted','rotation',270,'color','r','fontweight','bold','fontsize',13);

effort = [0 0 0];
% Determine the spatial prioritization
zz = num2str(prior);

if prior > 0 && prior < 1000 %sequential prioritization
% Figure out proportion of resources to each subpopulation
first = str2double(zz(1)); second = str2double(zz(2)); last = str2double(zz(3));
y1 = 1-(1/(1+alpha)^N(str2double(zz(1)))); % prioritization of resources to k1 given Nk1
y2p = 1-(1/(1+alpha)^N(str2double(zz(2)))); % prioritization function for k2 given Nk2 
y2 = (1-y1).*y2p; %proportion of remaining resources from k1 that go to k2 given fp(Nk2)
y3 = (1-y1)-y2; %proportion of remaining resources from k1 that go to k3 given fp(Nk2) and fp(Nk1)

elseif prior > 1000    
    if str2num(zz(2)) == 0 % if we are prioritizing one pop and then the other two equally
        first = str2double(zz(1)); second = str2double(zz(3)); last = str2double(zz(4));
        y1 = 1-(1/(1+alpha)^N(str2double(zz(1)))); % prioritization of resources to k1 given Nk1
        y2 = (1-y1)*0.5;
        y3 = y2;
    elseif str2num(zz(3)) == 0 % if we are prioritizing the first two pops equally and then prioritizing a third
        first = str2double(zz(1)); second = str2double(zz(2)); last = str2double(zz(4));
        y1 = (1-(1/(1+alpha)^(N(str2double(zz(1)))+N(str2double(zz(1))))))/2;
        y2 = y1;
        y3 = 1-(y1+y2);
    end
end
which = [y1 y2 y3];
%which = which./sum(which); % re-scale so that proportions sum to 1
temp = which.*beta; % determine actual intensity for each pop in this timestep based on monthly budget
effort = temp; % determine actual intensity for each pop in this timestep
effort(first) = temp(1);
effort(second) = temp(2);
effort(last) = temp(3);
end
```
