## Supplementary material for "Optimal spatial prioritization of control resources for elimination of invasive species under demographic uncertainty": Code for model: MetadataS1.docx

**Metadata S1**

Ecological Applications

We have provided three code files to show equations of the prioritization function and how the models were run. All code is provided as Matlab R2019b files.

Code S1. ‘culling3’ is a function that conducts the prioritization of control effort at each month.

Code S2. ‘MetapopulationModel_function3’ is the main function that conducts all processes over time for one set of parameters.

Code S3. ‘MetapopulationModel_iterate4’ is a script that defines multiple parameter sets for running the functions.
