## Supplementary material for "Optimal spatial prioritization of control resources for elimination of invasive species under demographic uncertainty": Code for model: MetapopulationModel_function3.m

```
% Population-level stochastic model 
%Nt = N0e^rt; r = log(Nt/N0)

% Define demographic parameters
% N0 = 1000;
% K = 1000;
% subpops = 3; %number of subpops
% years = 10;
% cscale = 0.5; % scaling factor on growth rate (this should = annual lambda)
% connect = 'InfiniteIslands';
% annualmig = 0.2;  % annual proportion that migrate from a donor pop to all recipients; 
% exogenous = [0 0 0]; % number of exogenous migrants each year in each subpop;
% curve = 0.03; % scaling parameter on density-dependent capture
% int = 1; % monthly proportion of population removed (MONTHLY UNITS OF EFFORT)
% prior = 123; % prioritization scheme
% price = 1000; % Price for max effort per month
% budget = 100000;
% savename = 'MetapopulationControl';

function [N,C,T,B,D,Parms] = MetapopulationModel_function3(N0,K,subpops,years,cscale,con,annualmig,alpha,...
    beta,prior,price,budget,dcost,damfunc,obs,obsV,alphaC)
time = 12*years;
CON = {'none','TwoIslandMainland','SteppingStone','TwoIslandMainlandReversed','ConnectedIslands','InfiniteIslands'};
connect = char(CON(con));
conc = [65    52    22    23    32    18    15    15     4    15     8    35]; %SREL data for conception frequency
c_time = conc./sum(conc); %monthly probability of conceiving (i.e., timing of births is stochastic)
c_time = c_time.*cscale; % seasonal trend in birth rates
r = repmat(c_time,1,years);

% Define matrices for different connectivity structures: rows = donor, cols
% = recipient

mig = annualmig/12;  %define migration rate for case without uncertainty 
  switch connect
    case 'none' % o o o
        conmat = [0 0 0; 0 0 0; 0 0 0];
    case 'TwoIslandMainland' % o <- o -> o
        conmat = [0 0 0; 0.5 0 0.5; 0 0 0];
    case 'SteppingStone' % o -> o -> o
        conmat = [0 1 0; 0 0 1; 0 0 0];
    case 'TwoIslandMainlandReversed' % o -> o <- o
        conmat = [0 1 0; 0 0 0; 0 1 0];
    case 'ConnectedIslands' % <-> o <- o -> o <-> 
        conmat = [0 0 1; 0.5 0 0.5; 1 0 0];
    case 'InfiniteIslands' % o <-> o <-> o <-> 
        conmat = [0 0.5 0.5; 0.5 0 0.5; 0.5 0.5 0];
  end

N = zeros(time,subpops); T = zeros(time,subpops); C = zeros(time,subpops); D = zeros(time,subpops); B = zeros(time,1);
N(1,:) = N0; B(1,:) = budget; D(1,:) = dcost;
for t = 2:max(time) 

    % TRUE ECOLOGICAL PROCESSES
    % BIRTHS IN MONTH t
    for s = 1:subpops
        N(t,s) = poissrnd(N(t-1,s) + r(t)*N(t-1,s)*(1 - (N(t-1,s)/K)),1);
    end
    
    % CONDUCT INTERNAL MIGRATION IN MONTH t
    Nmat = [N(t,1).*ones(1,subpops); N(t,2).*ones(1,subpops); N(t,3).*ones(1,subpops)];
    temp = binornd(Nmat,conmat.*mig); % these are the numbers that will move
    N(t,1) = max(0,N(t,1)-sum(temp(1,:))); N(t,2) = N(t,2)+temp(1,2); N(t,3) = N(t,3)+temp(1,3); % do the accounting
    N(t,2) = max(0,N(t,2)-sum(temp(2,:))); N(t,1) = N(t,1)+temp(2,1); N(t,3) = N(t,3)+temp(2,3);
    N(t,3) = max(0,N(t,3)-sum(temp(3,:))); N(t,1) = N(t,1)+temp(3,1); N(t,2) = N(t,2)+temp(3,2);
    
    % UNCERTAINTY FROM OBSERVATION OF N: assume we observe truth vs a
    % minimum
    
    switch obs
        case 0 % no error
            Nobs = N(t,:);
        case 1 % biased low
            pd = makedist('normal','mu',0.9,'sigma',obsV); pdT=truncate(pd,0,1); x = 0:0.01:1; yy=pdf(pdT,x); temp = randsample(x,3,'true',yy);
            Nobs = zeros(1,3);
            Nobs(1) = N(t,1)*temp(1); 
            Nobs(2) = N(t,2)*temp(2); 
            Nobs(3) = N(t,3)*temp(3);
        case 2 % biased high
            pd = makedist('normal','mu',0.9,'sigma',obsV); pdT=truncate(pd,0,1); x = 0:0.01:1; yy=pdf(pdT,x); temp = randsample(x,3,'true',yy);
            Nobs = zeros(1,3);
            Nobs(1) = N(t,1)*((1-temp(1))+1); 
            Nobs(2) = N(t,2)*((1-temp(2))+1); 
            Nobs(3) = N(t,3)*((1-temp(3))+1); 
        case 3 % error, no bias; same degree of error on either side of the truth
            pd = makedist('normal','mu',0.9,'sigma',obsV); pdT=truncate(pd,0,1); x = 0:0.01:1; yy=pdf(pdT,x); temp = randsample(x,3,'true',yy);
            Nobs = zeros(1,3);
            if N(t,1) > 0; Nobs(1) = randsample(N(t,1)*temp(1):N(t,1)*((1-temp(1))+1),1); else; Nobs(1) = 0; end
            if N(t,2) > 0; Nobs(2) = randsample(N(t,2)*temp(2):N(t,2)*((1-temp(2))+1),1); else; Nobs(2) = 0; end
            if N(t,3) > 0; Nobs(3) = randsample(N(t,3)*temp(3):N(t,3)*((1-temp(3))+1),1); else; Nobs(3) = 0; end
    end
    % CONDUCT CONTROL IN MONTH t  (NEED TO ADD PRIORITIZATION SCHEMES AND
    % EFFORT/COST RELATIONSHIP)
if sum(Nobs) > 0 && sum(beta) > 0
    prob = 1-(1./(1+alphaC).^Nobs); %density-dependent capture probability
    switch prior
        case 0            
            take = poissrnd(N(t,:).*prob.*beta/3); % stochastic take (prob = catchability based on N, beta = effort level)    
            C(t,:) = (beta/3).*price;
        otherwise
            effort = culling3(Nobs,prior,beta,alpha);
            take = poissrnd(N(t,:).*prob.*effort);
            if sum(Nobs) == 0; C(t,:) = 0; else; C(t,:) = effort.*price; end % assumes we are putting in full effort if there is > 1 pig
    end
            N(t,:) = max(0,N(t,:)-take);
            T(t,:) = take;
            B(t,1) = max(0,B(t-1,1)-nansum(C(t,:)));
            if B(t,1) == 0; beta(:) = 0; end % once we use all the budget; stop control
            if sum(N(t,:)) == 0; B(t) = NaN; end % stop drawing down budget if there are no more

    switch damfunc % damage cost function based on remaining pigs
        case 1 % saturating
            dd = 1-(1./(1+100).^min(1,N(t,:)./K)); % impact per subpopulation as a proportion of carrying capacity
        case 2 % linear
            dd = min(1,N(t,:)./K);             
        case 3 % exponential
            dd = 1-(1-(1./(1+100).^(1-min(1,N(t,:)./K))));
    end
    D(t,:) = dd.*dcost; % damage (0-1) as a function of population density
end
end
Parms = [N0,K,years,cscale,price,budget,alpha];

%save(savename,'N','C','T','B','Parms','ParmNames','-v7.3');

end
```
