## Supplementary material for "Optimal spatial prioritization of control resources for elimination of invasive species under demographic uncertainty": Code for model: MetapopulationModel_iterate4.m

```
% Iteration of main function for population-level stochastic model; 
% discrete time, monthly time step

% DEFINE FIXED PARAMETERS
N0 = 1000;
K = 1000;
subpops = 3; %number of subpops
years = 10; time = years*12;
savename = 'MetapopulationControl_U_082719_';
reps = 100; % repeated runs for the same parameter set
cscale = 0.5; % scaling factor on growth rate (this should = annual lambda)
% ParmNames = {'Initial N';'Carrying capacity';'Years for simulation';'Birth rate scaling';'Internal migration rate';'Exogenous migration rate';'Connectivity';...
%     'Control effort';'Prioritization of pops for control';'price per unit effort of control';'Total budget'};
dcost = 2000; % Maximum damage cost per month per subpop at carrying capacity 
price = 2000*subpops; %2000*subpops/int; % Price for max effort per month
budget = price*time;

%   switch connect
%     case 'none-1' % o o o
%         conmat = [0 0 0; 0 0 0; 0 0 0];
%     case 'TwoIslandMainland - 2' % o <- o -> o 
%         conmat = [0 0 0; 0.5 0 0.5; 0 0 0];
%     case 'SteppingStone - 3' % o -> o -> o
%         conmat = [0 1 0; 0 0 1; 0 0 0];
%     case 'TwoIslandMainlandReversed - 4' % o -> o <- o
%         conmat = [0 1 0; 0 0 0; 0 1 0];
%     case 'ConnectedIslands - 5' % <-> o <- o -> o <-> 
%         conmat = [0 0 1; 0.5 0 0.5; 1 0 0];
%     case 'InfiniteIslands - 6' % o <-> o <-> o <-> 
%         conmat = [0 0.5 0.5; 0.5 0 0.5; 0.5 0.5 0];
%   end
% DEFINE VARIABLE PARAMETERS FOR RUNS WITHOUT UNCERTAINTY
%%% Uncomment this section to run sims without uncertainty
% con = 1; % no connectivity    o o o
% prior = [0 123 1203 1023]; % prioritization scheme
% beta = [0.2 0.4 0.6 0.8]; % the max proportion by which the population can be reduced in one month (management intensity) 
% damf = 1:3;
% alpha = [0.0025 0.005 0.01 0.03]; %alpha_p
% alphaC = [0.013 0.03 0.07]; %alpha_c
% obs = 0; obsV = 0; uconStruc = 0; uconRate = 0;
% grid1 = allcomb(0,beta,prior,con,damf,obs,obsV,uconStruc,uconRate,alpha,alphaC);
% 
% con = 2; % TwoIslandMainland   o <- o -> o
% prior = [0 213 132 2013 1302]; % prioritization scheme
% annualmig = [0.01 0.025 0.05 0.075 0.1 0.125 0.15 0.2];
% grid2 = allcomb(annualmig,beta,prior,con,damf,obs,obsV,uconStruc,uconRate,alpha,alphaC);
% 
% con = 3; % SteppingStone  o -> o -> o
% prior = [0 123 321 1203 3201]; % prioritization scheme
% grid3 = allcomb(annualmig,beta,prior,con,damf,obs,obsV,uconStruc,uconRate,alpha,alphaC);
% 
% con = 4; % TwoIslandMainlandReversed  o -> o <- o
% prior = [0 132 213 1302 2013]; % prioritization scheme
% grid4 = allcomb(annualmig,beta,prior,con,damf,obs,obsV,uconStruc,uconRate,alpha,alphaC);
% 
% con = 5; % ConnectedIslands  <-> o <- o -> o <-> 
% prior = [0 213 132 2013 1302]; % prioritization scheme
% grid5 = allcomb(annualmig,beta,prior,con,damf,obs,obsV,uconStruc,uconRate,alpha,alphaC);
% 
% con = 6; % InfiniteIslands  o <-> o <-> o <-> 
% prior = [0 123 1203 1023]; % prioritization scheme
% grid6 = allcomb(annualmig,beta,prior,con,damf,obs,obsV,uconStruc,uconRate,alpha,alphaC);
% 
% grid = [grid1;grid2;grid3;grid4;grid5;grid6];
% id = find(grid(:,3) == 0 & grid(:,10) < 0.03); % varying alpha when prior = 0 is redundant so remove all those (was upposed to be 0.0025 but put 0.01 so now we have 3 of the same) 
% grid(id,:) = [];

% DEFINE VARIABLE PARAMETERS FOR RUNS WITH UNCERTAINTY
%%% Uncomment this section to run sims with uncertainty
con = 1; % no connectivity    o o o
prior = [0 1023 2013 3012 213 1203 1302]; % prioritization scheme (best = 1023) %TEST ALL OPTIMAL SCHEMES ON EACH TRUE CONNECTIVITY STRUCTURE
beta = [0.2 0.4 0.6 0.8]; % the max proportion by which the population can be reduced in one month (management intensity) 
damf = 2; % just run one
alpha = [0.0025 0.005 0.01 0.03]; %alpha_p
alphaC = 0.03; % just run one [0.013 0.03 0.07];
obs = 0; obsV = 0; uconStruc = 0; uconRate = 0;
%%% Uncomment this section to run sims without uncertainty
grid1 = allcomb(0,beta,prior,con,damf,obs,obsV,uconStruc,uconRate,alpha,alphaC);

con = 2; % TwoIslandMainland   o <- o -> o
prior = [0 1023 2013 3012 213 1203 1302]; %[0 213 132 2013 1302]; % prioritization scheme (best = 213)
annualmig = [0.01 0.025 0.05 0.075 0.1 0.125 0.15 0.2];
grid2 = allcomb(annualmig,beta,prior,con,damf,obs,obsV,uconStruc,uconRate,alpha,alphaC);

con = 3; % SteppingStone  o -> o -> o
prior = [0 1023 2013 3012 213 1203 1302];% [0 123 321 1203 3201]; % prioritization scheme (best = 1203)
grid3 = allcomb(annualmig,beta,prior,con,damf,obs,obsV,uconStruc,uconRate,alpha,alphaC);

con = 4; % TwoIslandMainlandReversed  o -> o <- o
prior = [0 1023 2013 3012 213 1203 1302]; %[0 132 213 1302 2013]; % prioritization scheme (best = 1302)
grid4 = allcomb(annualmig,beta,prior,con,damf,obs,obsV,uconStruc,uconRate,alpha,alphaC);

con = 5; % ConnectedIslands  <-> o <- o -> o <-> 
prior = [0 1023 2013 3012 213 1203 1302]; %[0 213 132 2013 1302]; % prioritization scheme (best = 213)
grid5 = allcomb(annualmig,beta,prior,con,damf,obs,obsV,uconStruc,uconRate,alpha,alphaC);

con = 6; % InfiniteIslands  o <-> o <-> o <-> 
prior = [0 1023 2013 3012 213 1203 1302]; %[0 123 1203 1023]; % prioritization scheme (best = 1023)
grid6 = allcomb(annualmig,beta,prior,con,damf,obs,obsV,uconStruc,uconRate,alpha,alphaC);

gridU = [grid1;grid2;grid3;grid4;grid5;grid6];

U = [0 0 0 0;...
    1 0.1 0 0;...
    1 0.25 0 0;...
    2 0.1 0 0;...
    2 0.25 0 0;...
    3 0.1 0 0;...
    3 0.25 0 0];
build = [];
for i = 1:size(U,1)
    build = [build; [gridU(:,1:5) repmat(U(i,:),size(gridU,1),1) gridU(:,10:11)]];
end

grid = build;
    
% Do the simulations
for jj = 1:reps
Budget = zeros(time,size(grid,1));
Ab = zeros(time,size(grid,1)); % abundance
Ta = zeros(time,size(grid,1)); % number removed
Rc = zeros(time,size(grid,1)); % removal costs
Dc = zeros(time,size(grid,1)); % damage costs
    for ii = 1:size(grid,1)
        vary = grid(ii,:);
        [N,C,T,B,D,~] = MetapopulationModel_function3(N0,K,subpops,years,cscale,vary(4),vary(1),vary(10),...
            vary(2),vary(3),price,budget,dcost,vary(5),vary(6),vary(7),vary(11));
        summary = [N T C D B];
        Budget(:,ii) = B;
        Ab(:,ii) = sum(N,2);
        Ta(:,ii) = sum(T,2);
        Rc(:,ii) = sum(C,2);
        Dc(:,ii) = sum(D,2);
        fprintf('Parameter Set: %d\n',ii)        
    end
     fprintf('Rep: %d\n',jj)      
     save(sprintf('%s%d',savename,jj),'Ta','Ab','Rc','Dc','Budget','grid','-v7.3');
end    
```
